# Prolonged HPA axis dysregulation in postpartum depression associated with adverse life experiences during development: a cross-species translational study for a novel therapeutic approach

**DOI:** 10.1101/2021.10.13.464258

**Authors:** Minae Niwa, Sedona Lockhart, Daniel J. Wood, Kun Yang, Jose Francis-Oliveira, Kyohei Kin, Adeel Ahmed, Gary S. Wand, Shin-ichi Kano, Jennifer L. Payne, Akira Sawa

## Abstract

Stress during childhood and adolescence increases the risk for postpartum depression (PPD). Patients with depression who have experienced adverse life events tend to be treatment refractory. However, the mechanism by which stress during childhood and adolescence are involved in the pathophysiology of PPD remains unclear. We investigated the longitudinal effects of adolescent stress on the hypothalamic-pituitary-adrenal (HPA) axis and behaviors in the postpartum period through mouse and human studies. We observed that adolescent social isolation caused an aberrantly sustained elevation of glucocorticoids via dysregulation of the HPA axis, leading to long-lasting postpartum behavioral changes in female mice. The postpartum behavioral changes elicited by this adolescent stress were not ameliorated by the medicines currently used for PPD treatment. However, a post-delivery treatment with a glucocorticoid receptor antagonist effectively ameliorated the behavioral changes in mice. We also demonstrated a significant impact of stress during childhood and adolescence on the HPA axis dysregulation and PPD in women. We provide experimental evidence that suggests a mechanism-driven therapeutic strategy (repurposing a GR antagonist) for at least some cases of treatment refractory PPD.

## Introduction

Pregnancy and delivery are events accompanied by significant physical and psychological changes to the mother ^1^. Mood disturbances and cognitive impairments affecting mothers during the postpartum period are common and lead to serious mental health problems, which can consequently affect the child’s development and behavior ^2,3^. Adverse life events, such as poor family/social support or a history of psychiatric disorders, are known as predominant risk factors for postpartum depression (PPD) ^1,2,4,5^. Women who experienced adverse life events are three times more likely to have PPD than women who did not experience any adverse life events ^6^. Furthermore, patients with depression who experienced adverse life events tend to be treatment refractory ^7,8^.

Mothers display endocrinological changes during pregnancy and the postpartum period. Such dynamic changes are observed in lactogenic hormones, including prolactin and oxytocin ^9^. The hypothalamic-pituitary-adrenal (HPA) axis is also activated during pregnancy and delivery, which results in an increase of glucocorticoid production ^10–12^. Furthermore, levels of estrogen, progesterone, and allopregnanolone are increased during pregnancy and followed by a precipitous drop-off after delivery ^4,13–16^. Thus, the rapid decline in reproductive hormones immediately after delivery is believed to participate in the onset of PPD ^4,17^. Consistent with this notion, postpartum injections of estrogen to women with PPD after delivery have shown some beneficial effects ^18–21^. Meanwhile, a sudden withdrawal from a two-month exposure of estradiol and progesterone at supra-physiological doses can induce depressive symptoms in women who had a history of PPD ^19^. In line with these human data, a postpartum withdrawal from exogenous estrogen and progesterone injections in rodents lead to behavioral changes in the forced swim test (FST), which has been frequently employed to evaluate many antidepressants including those that are currently used for PPD ^22–24^. Thus, this rodent model has shed light on the biological and behavioral changes associated with the rapid decline in reproductive hormones after delivery. However, these observations have not answered a crucial medical question of how exposures to early adverse life events underlie the pre-symptomatic pathological progression and the final emergence of postpartum behavioral changes relevant to PPD. Development of relevant animal models is warranted to fill the knowledge gap.

The first-line pharmacological treatment for PPD is selective serotonin reuptake inhibitors (SSRIs), particularly sertraline due to its very low breastmilk transmission to infants ^25^. Although these are effective compared to placebo, only about 54% of patients respond according to a 2015 report ^26^, indicating a need for novel pharmacological treatments. Another limitation is that symptomatic relief only appears several weeks after the initial medication ^27,28^. To overcome these limitations, an intravenous formulation of allopregnanolone (brexanolone), a positive allosteric modulator of GABA_A_ receptors (GABA_A_R), has been introduced to the repertoire of treatment for PPD ^16^. However, its drawbacks include cost and treatment accessibility (e.g., high price, intravenous administration, requirement of a Risk Evaluation and Mitigation Strategy, need for hospitalization) ^16,28–30^. Another concern is a potential temporary separation from the infant during a critical period of mother-infant bonding ^30^. Thus, it is still important to investigate other treatment avenues for PPD. A mechanistic understanding of how adverse life experiences during development, such as adolescent stress, influences the disease pathophysiology may provide clues for novel treatment approaches.

## Results

### Long-lasting changes of postpartum behaviors in dams exposed to adolescent social isolation

Adolescence is a special period when hormonal levels change dramatically and neural circuits are fine-tuned ^31^. Social isolation has been widely recognized as a significant stressor in the developmental period in mice ^32–36^. Our group has studied the influences of social isolation in adolescence on adult behaviors, including our recent report that suggests the utility of this paradigm in detecting postpartum cognitive changes in mice ^37–41^. In the present study, applying this paradigm, we examined the effects of adolescent psychosocial stress on PPD-related neuroendocrinological and behavioral changes in dams.

Experimental groups consisted of a) unstressed virgins, b) stressed virgins, c) unstressed dams and d) stressed dams (i.e. mice that were exposed to adolescent social isolation and gave the first-time birth to pups in adulthood) (**Extended Data Figure 1**). We chose the tail suspension test (TST) and FST as major assays, because their frequent use in discovering and evaluating many antidepressants including those that are currently used for PPD: although it is very difficult to capture several features in human depression (face validity) by mouse behavioral assays, the utility of these tests that evaluate a switch from active to passive behavior upon an acute stress, underlying behavioral adaptation and survival, has been proven in pharmacological studies for depression ^42^. Indeed, in the drug discovery process for PPD, FST was used for the discovery of an allopregnanolone analog, GABA_A_R modulator ^43^. We did not observe any differences in the TST and FST among the four groups at postpartum days 0 and 1, respectively (**Figure 1**). Dams exposed to adolescent social isolation (stressed dams), compared with other three groups, showed increased immobility time during the TST and FST at postpartum days 7 and 8, respectively (**Figure 1**). We also performed the sucrose preference test (SPT) to assess hedonic responses to sweet solutions ^44^: stressed dams showed a decrease in sucrose preference, starting on postpartum day 7, compared to unstressed dams (**Figure 2**). The behavioral changes in stressed dams in TST, FST, and SPT were prolonged for at least three weeks after delivery (**Figures 1** and **2**).

**Figure 1.**
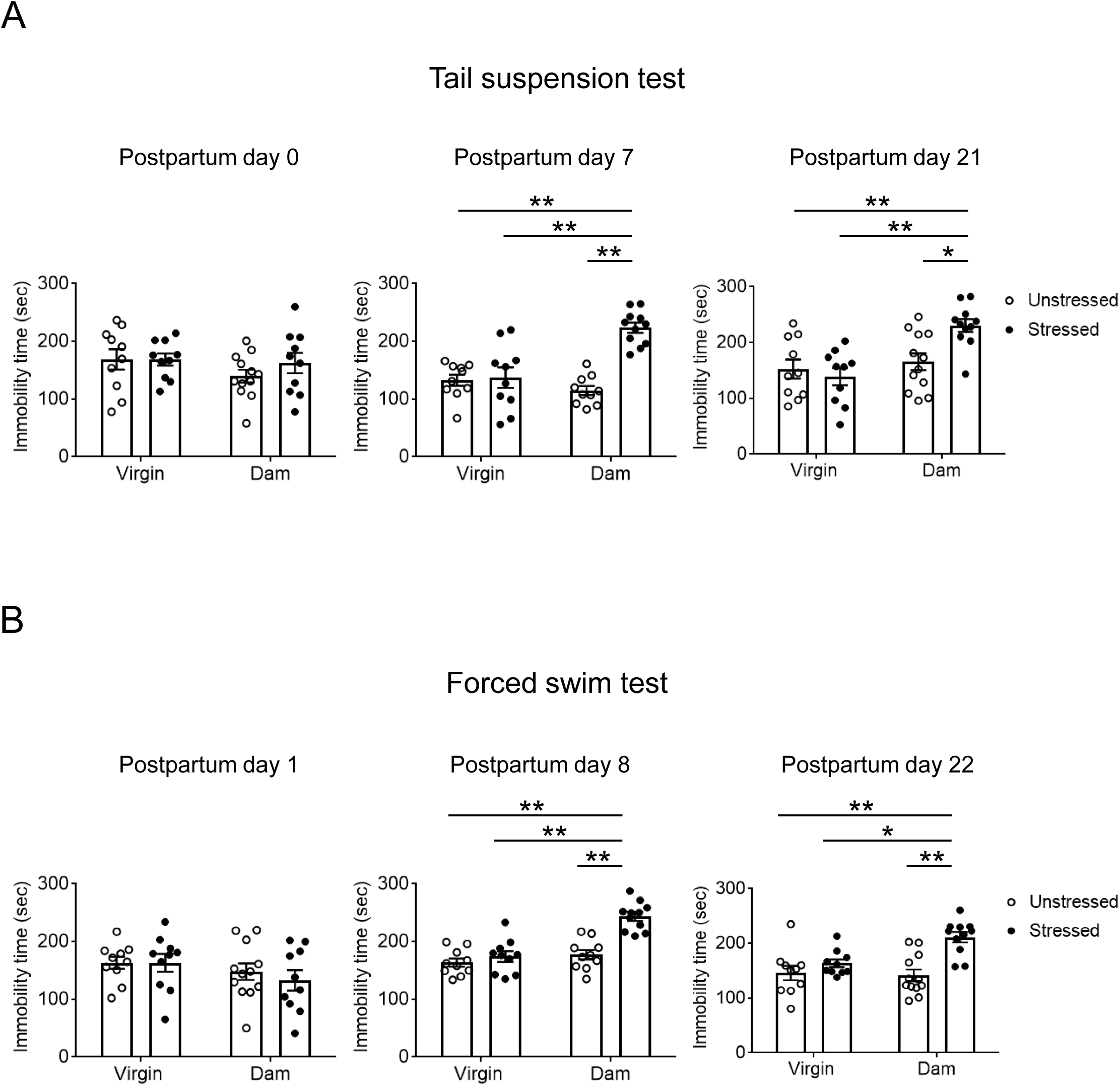
Long-lasting behavioral changes in the tail suspension and forced swim tests in dams exposed to adolescent social isolation. **A**, **B**, Immobility time (seconds) during the tail suspension (**A**) and forced swim tests (**B**) were assessed at postpartum days 0 and 1, postpartum days 7 and 8, and postpartum days 21 and 22, respectively. Behavioral changes emerged at 1 week postpartum and remained until at least 3 weeks postpartum. No changes across groups were observed immediately after delivery. N=10-12. Values are represented as mean ± SEM; \*\**P*<0.01 and \**P*<0.05.

**Figure 2.**
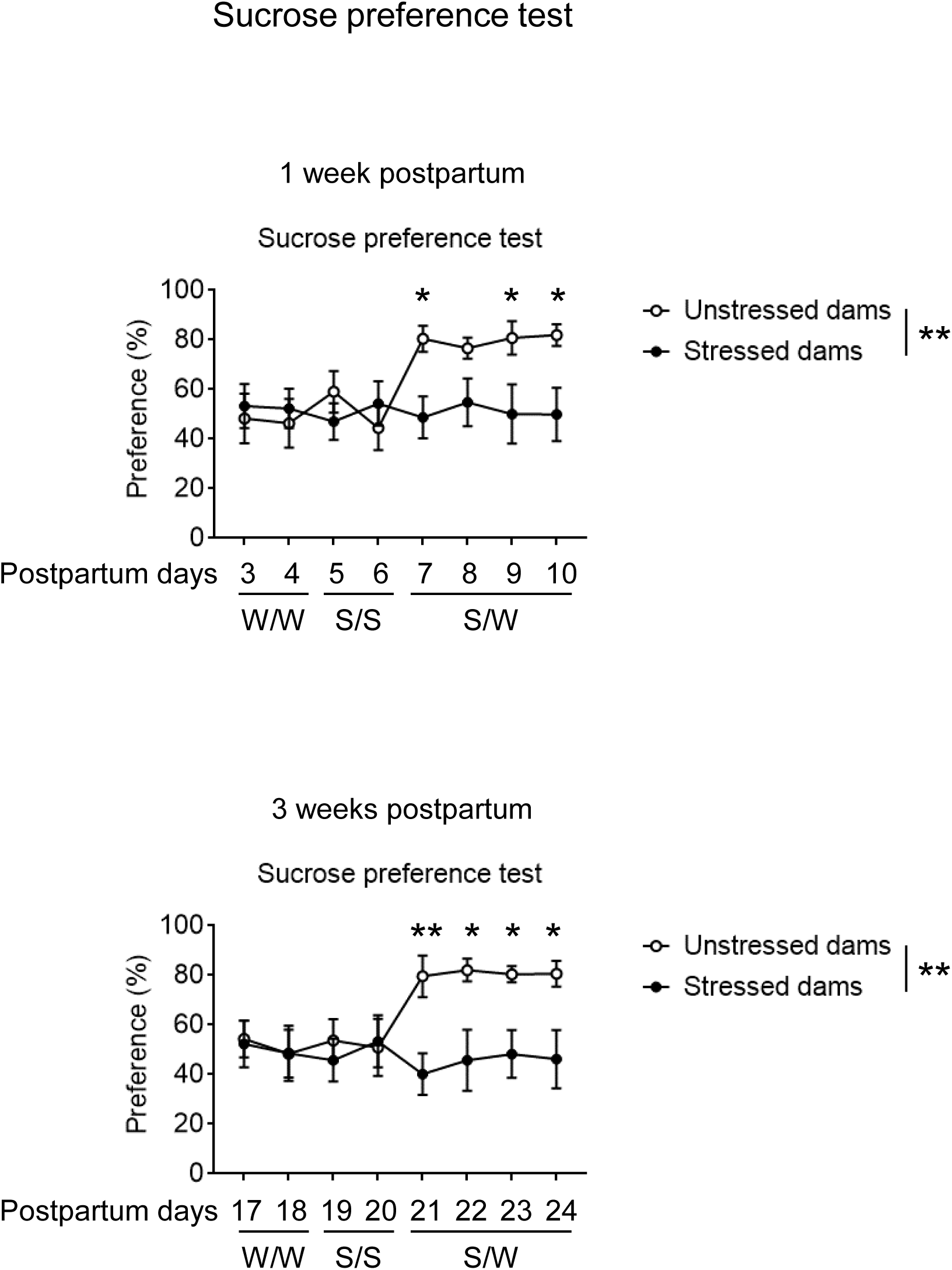
Long-lasting behavioral changes in the sucrose preference test in dams exposed to adolescent social isolation. Sucrose preference was measured at one-week (postpartum days 7-10) and three-week postpartum (postpartum days 21-24). Stressed dams showed a reduction in sucrose preference when given a choice between 1.5 % sucrose and water. N=11. W/W, water/water. S/S, 1.5 % sucrose/1.5 % sucrose. S/W, 1.5 % sucrose/water. Values are represented as mean ± SEM; \*\**P*<0.01 and \**P*<0.05.

Social cognition in mothers may be regulated to monitor and interpret social signals from others, and safely navigate the new living environment ^45^. Thus, we examined the effect of adverse adolescent stress on dams’ postpartum behavior in the three-chamber social interaction test (SIT) which is used to test the social communication ability in rodents ^46^. No behavioral differences among the groups were observed at postpartum day 0 in both sociability and social novelty recognition in the SIT (**Extended Data Figure 2**). At postpartum days 7 and 21, stressed dams, compared with unstressed virgins, showed a significant difference in social novelty recognition, but not in sociability (**Extended Data Figure 2**). These data suggest that adolescent stress, combined with the events of pregnancy and delivery, leads to prolonged changes in postpartum behaviors in TST, FST, SPT, and SIT in dams.

### Prolonged elevation of corticosterone levels and altered glucocorticoid signaling in dams exposed to adolescent social isolation: its pivotal role in long-lasting changes of postpartum behaviors

Many studies have reported alterations in plasma hormone levels in patients with PPD and a correlation between hormonal changes and onset of PPD symptoms ^9,19–22,47–49^. Thus, we examined the plasma levels of estradiol, progesterone, prolactin, oxytocin, and corticosterone in our animal model. Although we observed physiological changes in the levels of estradiol, progesterone, prolactin, and oxytocin associated with pregnancy and delivery, no further differences in these changes were observed between unstressed and stressed dams at any time point (**Extended Data Figure 3**). In contrast, the level of plasma corticosterone in both unstressed and stressed dams was increased after late pregnancy in comparison to virgin mice (**Figure 3A**). Corticosterone levels in unstressed dams began to decrease after delivery (**Figure 3A**). Notably, corticosterone levels in stressed dams were consistently higher than those in unstressed controls at 1 and 3 weeks postpartum (**Figure 3A**). These data suggest that a potential change in the control of the HPA axis may occur in stressed dams, resulting in a prolonged elevation of corticosterone levels. Positive correlations between the levels of plasma corticosterone and behavioral changes in the TST and FST were observed at 1 week and 3 weeks postpartum (**Extended Data Figure 4**).

**Figure 3.**
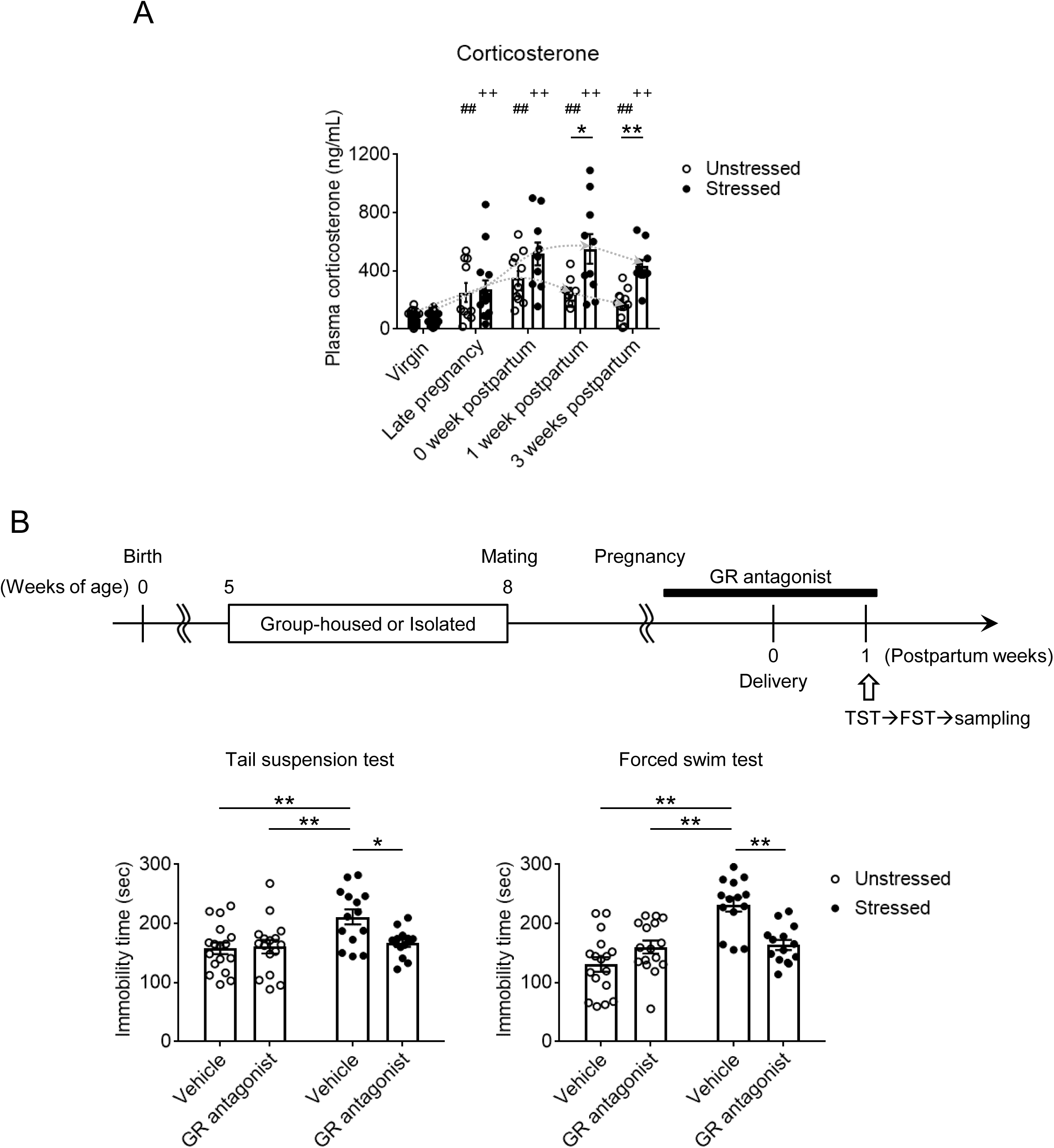
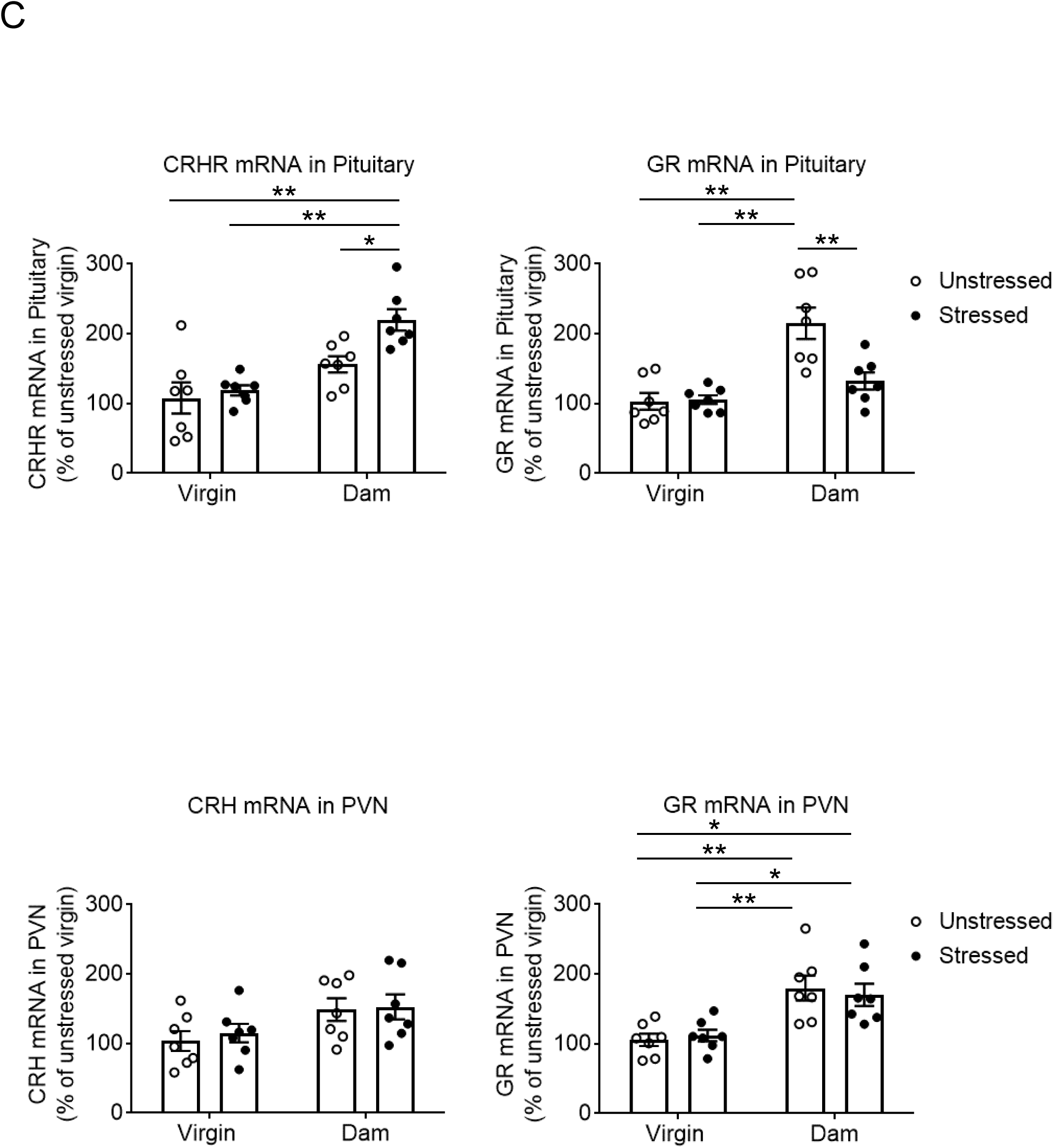
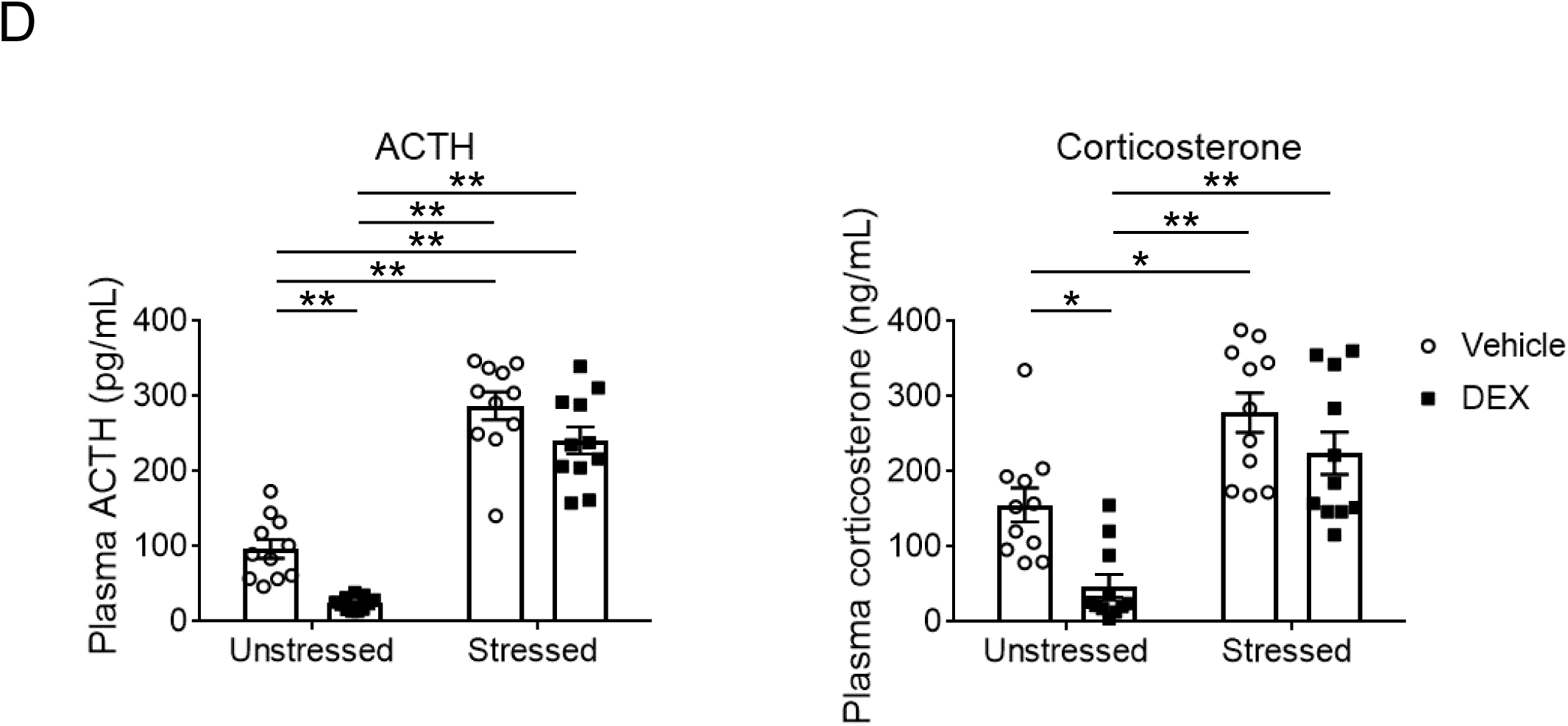
Prolonged elevation of corticosterone levels and altered glucocorticoid signaling in dams exposed to adolescent social isolation: its pivotal role in long-lasting changes of postpartum behaviors. **A**, Levels of corticosterone were measured at five time points: virgin, late pregnancy, 0 week postpartum, 1 week postpartum, and 3 weeks postpartum. Corticosterone levels in stressed dams were consistently higher than those in unstressed controls at 1 and 3 weeks postpartum. N=9-30. **B**, Causal effects of the GR on postpartum behavioral changes in the TST and FST. CORT113176 (80 mg/kg, *p.o.*, once daily from gestation day 14 to 24 h prior to sampling at postpartum day 9) ameliorated the behavioral changes in the tail suspension and forced swim tests in stressed dams at postpartum days 7 and 8, respectively. The antagonist did not affect behavior in the TST and FST in unstressed dams. N=14-17. **C**, Differences in CRHR and GR mRNA levels in the pituitary, but not PVN, between unstressed and stressed dams. Differences in the mRNA levels of CRHR and GR in the pituitary were observed between unstressed and stressed dams at postpartum day 9. No difference in the mRNA levels of CRH and GR in the PVN was observed between unstressed and stressed dams at postpartum day 9. There was no difference in the mRNA levels examined in this study between unstressed and stressed virgin mice. N=7. **D**, Changes in HPA axis negative feedback in stressed postpartum dams. In the dexamethasone (DEX) suppression test, DEX (0.1 mg/kg, *i.p.*) was administered at postpartum day 8. Treatment with low dose DEX reduced the levels of ACTH and corticosterone in unstressed dams, whereas it failed to suppress these hormones in stressed dams. N=11. Values are represented as mean ± SEM; \*\**P*<0.01 and \**P*<0.05; ^##^*P*<0.01 versus unstressed virgins; ^++^*P*<0.01 versus stressed virgins.

The above results suggest that the alteration in the HPA axis may underlie the behavioral changes observed in stressed dams. To address a role of the altered glucocorticoid signaling in behavioral changes during the postpartum period, a selective GR antagonist, CORT113176, was orally administered from gestation day 14 to 24 h prior to sampling at postpartum day 9, one day after the last behavioral testing at one-week postpartum when we first observed significant behavioral changes in stressed dams. The CORT113176 administration blocked the behavioral changes in the TST and FST in stressed dams at postpartum days 7 and 8, respectively (**Figure 3B**). These results support the hypothesis that prolonged activation of glucocorticoid signaling underlies long-lasting behavioral changes in dams exposed to adolescent social isolation.

### Molecular expression changes in the HPA axis in dams exposed to adolescent social isolation

In the HPA axis, corticotropin releasing hormone (CRH) from the paraventricular nucleus (PVN) of the hypothalamus triggers the secretion of adrenocorticotropic hormone (ACTH) from the anterior pituitary gland, leading to the production of glucocorticoids from the adrenal cortex, which in turn regulate GRs on the PVN and pituitary gland by a negative feedback mechanism^50,51^.

We hypothesized that mice exposed to adolescent social isolation might show molecular expression changes in the HPA axis after delivery, compared to mice without this adolescent social isolation. To address this question, we measured the mRNA levels of CRH and GR in the PVN, and those of CRH receptor1 (CRHR) and GR in the pituitary gland at 1 week postpartum when stressed dams start to show postpartum behavioral changes described above. We compared the expression data among stressed dams, unstressed dams, age-matched stressed virgins, and age-matched unstressed virgins. Delivery itself augmented the expression of GR in both the PVN and pituitary gland in unstressed mice (**Figure 3C**). In stressed dams, this delivery-associated augmentation of GR was not observed in the pituitary whereas it was still seen in the PVN (**Figure 3C**). Regarding the CRH-CRHR, we observed an upregulation of CRHR in the pituitary gland only in stressed dams (**Figure 3C**). The changes seen in the pituitary gland of stressed dams indicated that adolescent social isolation might affect at least the pituitary gland after delivery.

To validate this notion at the functional levels, we performed a low-dose dexamethasone (DEX) suppression test ^52,53^ at 1 week postpartum. DEX, a synthetic glucocorticoid, does not effectively penetrate the blood-brain barrier (BBB) when used at low doses (e.g., 0.1 mg/kg) ^52–54^. The low dose of DEX binds to GR in the anterior pituitary gland, but not the PVN, because the anterior pituitary is located outside the BBB ^54–56^. Administration of a low dose (0.1 mg/kg) of DEX reduced the levels of ACTH and corticosterone in unstressed dams, whereas it failed to suppress these hormones in stressed dams (**Figure 3D**).

### Normalization of postpartum behavior changes in dams exposed to adolescent social isolation following a post-delivery treatment with a GR antagonist

It is possible to hypothesize that the aforementioned mouse model may represent an important subset of PPD patients who experience adverse life events and resultant changes in the HPA axis. Because half of patients with PPD are refractory to SSRI treatment and these treatment refractory cases are likely to be associated with adverse life events ^7,8^, we investigated the utility of our adolescent stress paradigm in modeling these clinically difficult cases. To this end, we examined the beneficial effects of post-delivery treatment with the SSRI, the GABA_A_R modulator, or the GR antagonist. Treatment with an SSRI (fluoxetine) at 18, 36, and 54 mg/kg for one-week post-delivery did not normalize the behavioral changes in TST and FST in stressed dams (**Figure 4** and **Extended Data Figure 5**). Treatment of the same duration with a GABA_A_R modulator (ganaxolone) at 10, 20, and 30 mg/kg was also ineffective (**Figure 4** and **Extended Data Figure 5**). Post-delivery treatment for one-week with a GR antagonist (CORT113176) at 40 mg/kg ameliorated the behavioral changes only in TST in stressed dams, while an 80 mg/kg dose normalized the behavioral changes in both the TST and FST (**Figure 4** and **Extended Data Figure 5**). Treatment with the compound at 160 mg/kg had no effect in both TST and FST in stressed dams **(Extended Data Figure 5)**.

**Figure 4.**
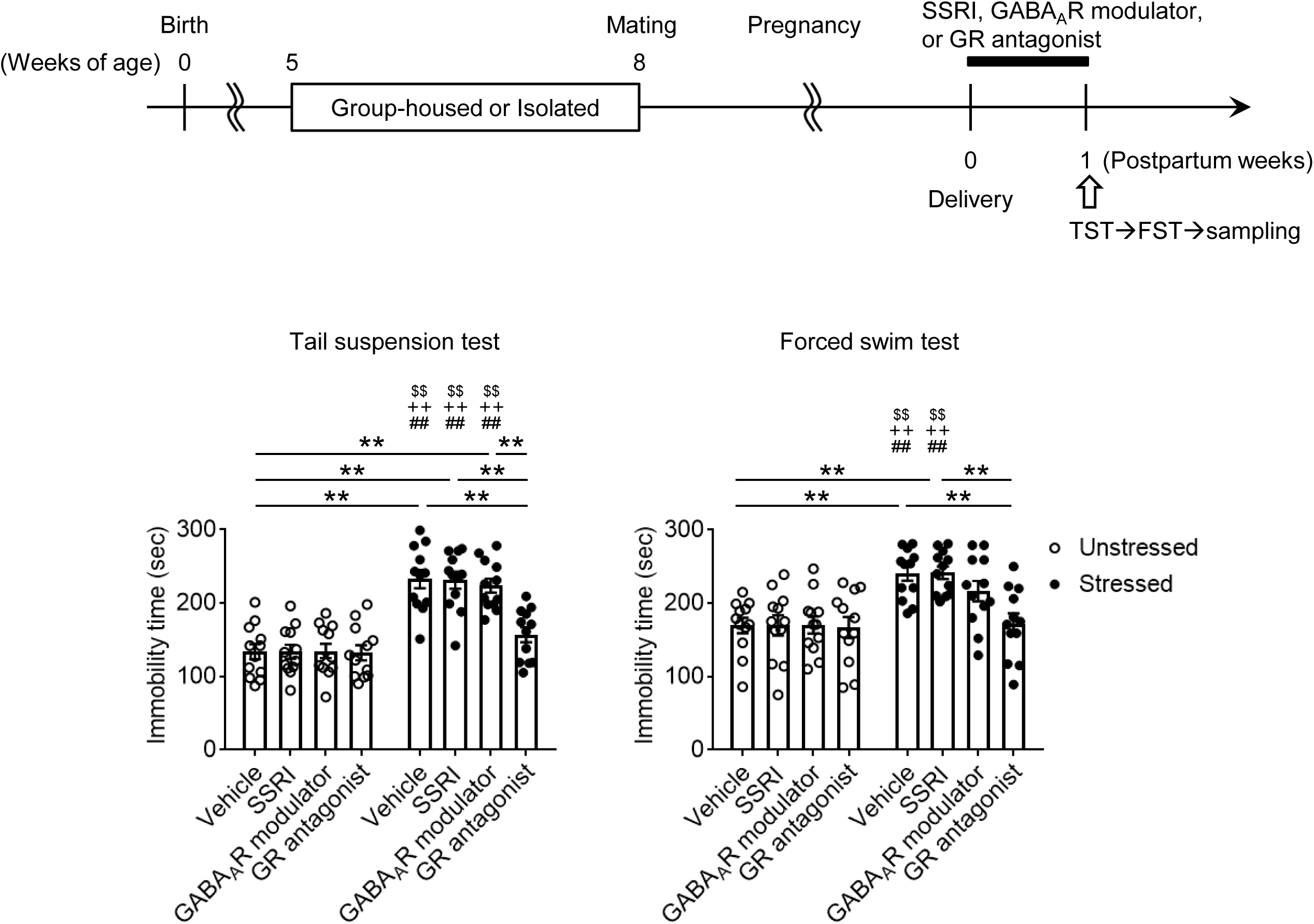
Normalization of postpartum behavior changes in dams exposed to adolescent social isolation following a post-delivery treatment with a GR antagonist. Mice were treated with an SSRI fluoxetine (18 mg/kg, *p.o.*), a GABA_A_ receptor modulator ganaxolone (10 mg/kg, *i.p.*), or a GR antagonist CORT113176 (80 mg/kg, *p.o.*) once daily from postpartum day 0 to 24 h prior to sampling at postpartum day 9. Post-delivery treatment with only the GR antagonist ameliorated the behavioral changes in the tail suspension and forced swim tests in stressed dams. N=12. Values are represented as mean ± SEM; \*\**P*<0.01; ^##^*P*<0.01 versus unstressed dams treated with the SSRI; ^++^*P*<0.01 versus unstressed dams treated with the GABA_A_ receptor modulator; ^$$^*P*<0.01 versus unstressed dams treated with the GR antagonist.

### A link between early life stress, a sustained increase in the glucocorticoid signaling, and PPD in humans

To examine whether our preclinical model shows clinically-relevant biology and behavioral phenotypes, we assessed the longitudinal influence of stress during childhood and adolescence on HPA axis function and postpartum mental conditions in humans. Specifically, we examined the relationship between stress during childhood and adolescence [a history of major mental illness, abnormal home environment, abnormal childhood behavior, and/or traumatic events], the HPA axis, and the development of PPD in 116 women. We observed increased risks for developing PPD by stress, in particular a history of mental illness (**Figures 5A** and **5B**). We then compared the level of plasma cortisol, as a surrogate of HPA axis activity, between non-PPD women who had a previous mental illness diagnosis and PPD patients who had a previous mental illness diagnosis. The former group showed an equivalent peak of glucocorticoids leading up to delivery followed by a gradual decline (**Figure 5C**). In contrast, cortisol levels in women with PPD who had a previous mental illness diagnosis exhibited both elevated and sustained levels of plasma cortisol until at least six weeks postpartum (**Figure 5C**).

**Figure 5.**
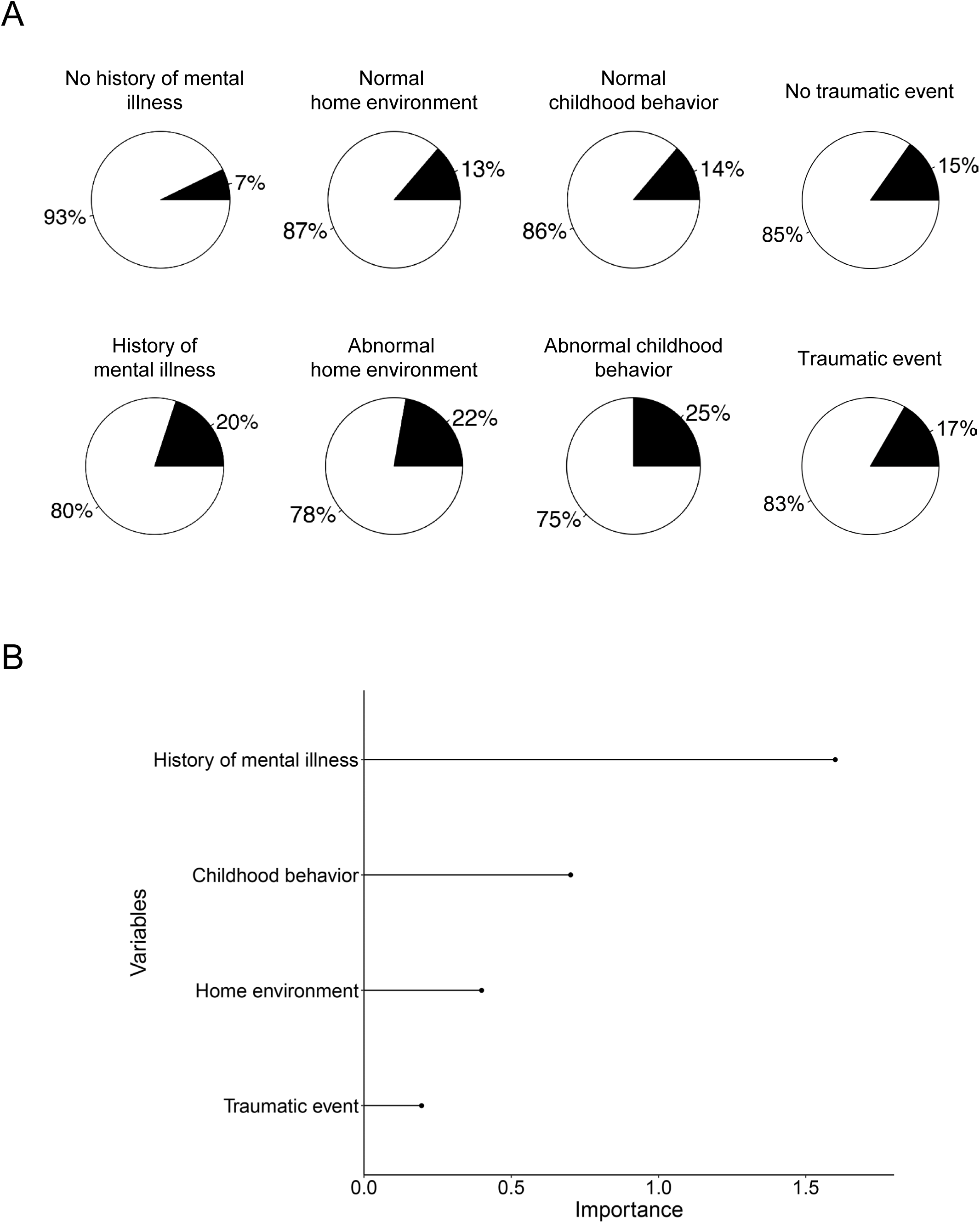
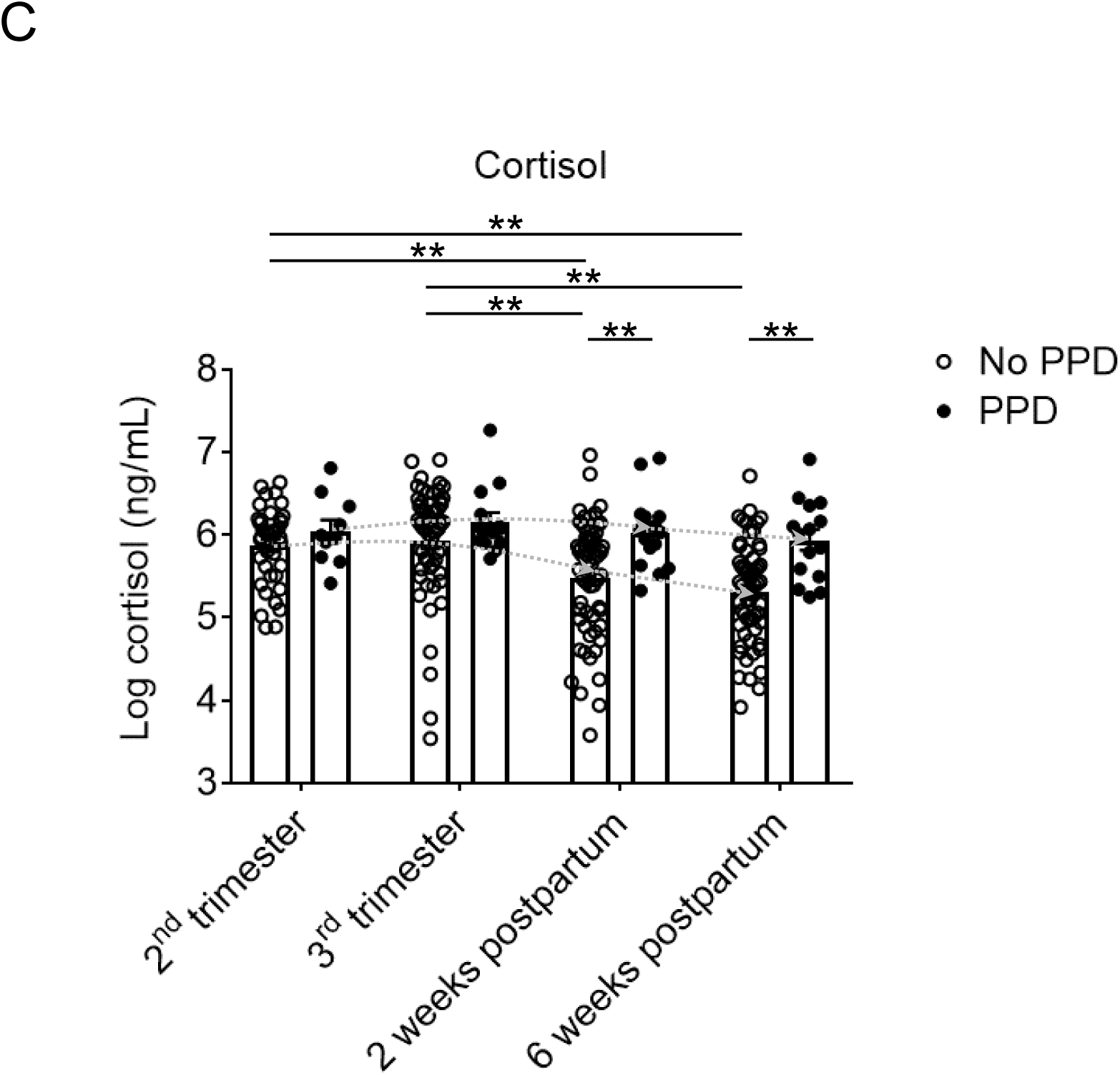
A link between early life stress, a sustained increase in the glucocorticoid signaling, and PPD in humans. **A**, Significance of adverse early life events on PPD in human subjects. Participants with a history of mental illness were more likely to be diagnosed with PPD than those with other risk factors. The white and black slices of the pie charts show the percentages of participants without and with PPD, respectively. N=116. **B**, Risk factors for PPD in human subjects. A history of mental illness was the most important risk factor positively correlated and linked to the development of PPD compared to childhood behavior, home environment, and traumatic events. N=116. **C**, A prolonged elevation in plasma cortisol levels in PPD patients with a history of mental illness. Participants with a history of mental illness and no diagnosis of PPD showed a significant decline in cortisol levels after delivery. Participants with a history of mental illness and PPD showed a sustained elevation of cortisol levels until at least 6 weeks postpartum. N=10-60. Values are represented as mean ± SEM; \*\**P*<0.01 and \**P*<0.05.

## Discussion

The main findings from the present study include that an adolescent social isolation elicits a prolonged elevation of corticosterone levels and glucocorticoid signaling in mice, which in turn results in long-lasting postpartum behavioral changes. In parallel, we assessed the longitudinal influence of stress during childhood and adolescence on HPA axis function and postpartum mental conditions in humans and supported the notion that the data from our mouse model implies clinically-relevant biology and behavioral phenotypes. We believe that the discovery in the present study sheds light on the relationship between adolescent stress and postpartum behavioral changes, bridging a major knowledge gap in the understanding of PPD.

We pharmacologically proved that the alteration in the glucocorticoid signaling after delivery accounted for the behavioral changes. Of the most importance, we demonstrated that a short-term (one-week), post-delivery treatment, against this pathological mechanism is sufficient to block the behavioral changes in stressed dams, whereas other medicines currently used for PPD in the clinical setting are ineffective. Furthermore, we observed a sustained elevation of cortisol in patients with PPD with a history of mental illness. Together, we suggest that adolescent stress can underlie prolonged behavioral abnormalities in the postpartum period, likely mediated by long-lasting changes in the HPA axis.

PPD is a “specifier” in Depressive Disorders in the Diagnostic Statistical Manual (DSM). This distinction helps tailor treatment to specific triggers and contexts, which is analogous to the application of precision medicine in oncology^57–59^. The novelty of the present study lies in discovering and defining the role and mechanism of the HPA axis and glucocorticoid signaling in the context of PPD. In particular, different from previous studies on PPD, we underscore adolescent psychosocial stress and its longitudinal impact on the disease after delivery.

The mechanisms of how GABA_A_R modulators exert beneficial influences on PPD are still actively being studied ^60–63^. Intervention with GABA_A_R-mediated neurotransmission in the PVN may be one of these mechanisms ^60^. Under our experimental conditions, in which we selected the doses and administration routes of the compounds based on published protocols ^64–67^, neither a GABA_A_R modulator nor an SSRI ameliorated the behavioral changes. We believe that our findings do not diminish the value of current drugs for PPD. Instead, our present study indicated that the beneficial effect of the post-delivery GR antagonist administration appeared quickly (only within one week postpartum). We hope that the finding has the potential to reshape the current approaches to PPD treatment, in particular for the cases with adverse life events in adolescence.

The systemic administration of the medicines we conducted in the present study may possibly elicit confounding factors, in particular those associated with their impact on metabolic pathways ^68,69^. However, at least in our experimental conditions, none of the medicines, including the GR antagonist, affected body weight (**Extended Data Figure 6**). We also confirmed that the GR antagonist had no effect on blood glucose levels and weight of the thymus, spleen, and visceral fat (**Extended Data Figure 7**). These data suggest that the influences of these medicines on body metabolism may be minimal.

The present data indicate molecular changes at least in the anterior pituitary gland in the HPA axis in our PPD model. Given that there were no differences in the levels of oxytocin and prolactin (released by the posterior lobe) between unstressed and stressed dams, it is unlikely that the posterior pituitary gland is affected in the present model. Further studies with region- or circuit-specific manipulations of the GR and related molecules will facilitate an understanding of the circuitry mechanisms that are sensitive to adolescent stress and their impact on HPA axis regulation and postpartum behaviors. There are specific brain regions that have been associated with adolescent stress ^50,70^. For example, in a recent publication from our group ^41^, we suggested a role of GR in the anterior insula (AI)-prelimbic cortex (PrL) pathway in altered behaviors in stressed dams during the postpartum period. Given that the AI and PrL are sensitive to stressors ^71–74^, it is an interesting question of how changes in this circuit influence the HPA axis underlying postpartum behaviors, particularly by linking to epigenetic mechanisms underlying the prolonged changes in the HPA axis and subsequent behaviors.

Previous studies on PPD have frequently examined the roles of reproductive steroid hormones ^4,13–16^. Although the goal of preclinical studies is usually to shed light on novel mechanisms, the success of subsequent translational studies will be supported when multifaceted information from preclinical studies is well integrated. Therefore, to extrapolate this preclinical observation to human studies, more frequent sampling of blood and examination of glucocorticoid signaling, together with other reproductive steroid hormones, in greater detail will be beneficial. Although mouse behaviors are useful outcome measures to understand the impact of disease pathophysiology, we would advocate that clinical trials will be the only and final way of drawing the conclusions regarding the therapeutic potential of GR antagonists for human PPD, particularly the cases with adverse life events during development.

Although other environmental stressors or complex genetic factors may also underlie PPD, the goal of the present study is to establish a straightforward model with a single environmental stressor under a homogenous genetic condition, which allowed us to dissect a role of an altered glucocorticoid signaling in long-lasting behavioral changes in the postpartum period. Mouse models are regarded as useful in translational research because their genetic homogeneity, in contrast to the heterogeneity of human populations, leads to less variability in experimental results ^75,76^. A mild social isolation in adolescence is known to cause consistent behavioral changes in genetically vulnerable mice or dams, particularly the inbred mouse stains we used here ^37–41^. Future studies with this unique model may include studies of maternal behaviors, transgenerational transmission of risk to children reared by stressed mothers, involvement of the mineralocorticoid receptor, differences between depression during pregnancy and PPD, and involvement of genetics and epigenetics in the pathological mechanisms underlying PPD.

Here, we underscored the potential of a GR antagonist for the treatment of a subset of PPD. GR antagonists can be orally administered in treatment of Cushing syndrome (mifepristone) as well as in other ongoing clinical trials ^77,78^. A post-delivery, short-term treatment of a GR antagonist effectively normalizes postpartum behavioral changes in our mouse model, whereas medicines currently used for PPD treatment were ineffective within this condition. Accordingly, our data would support the notion that repurposing a GR antagonist may lead to an immediate improvement for at least some cases of treatment refractory PPD.

## Methods

### Animals

To determine the effects of adolescent psychosocial stress on postpartum behaviors in adulthood, healthy virgin C57BL/6J female mice were exposed to mild isolation stress during late adolescence (from 5 to 8 weeks of age), which alone caused no endocrine or behavioral changes ^37^. Each mouse was then mated with a C57BL/6J male mouse at 8 weeks of age and gave birth to pups. This group was designated “stressed postpartum mice”. Isolation consisted of no interaction with other mice and confinement to opaque wire-topped polypropylene cages, whereas group-housed mice were kept in clear wire-topped plastic cages (21×32×13 cm). Unstressed virgins, stressed virgins, unstressed dams, and stressed dams were age-matched in all the experiments. All mice were maintained under a controlled environment (23 ± 3°C; 40 ± 5% humidity; light and dark cycles started at 7 am and 9 pm, respectively) with ad libitum access to food and water. All experimental procedures were performed in accordance with the National Institutes of Health Guidelines for the Care and Use of Laboratory Animals, under the animal protocols approved by the Institutional Animal Care and Use Committees at the Johns Hopkins University and the University of Alabama at Birmingham.

### Drug Treatments in mice

To assess the influence of glucocorticoid signaling on behavioral changes related to mobility/despair and social cognition in stressed postpartum mice, the selective GR antagonist, CORT113176 [80 mg/kg, *p.o*, (oral gavage with flexible polypropylene feeding tubes) once a day, dissolved in dimethyl sulfoxide and further dissolved in saline containing 0.5 % hydroxypropyl methylcellulose + 0.1 % Tween 80], was orally administered from gestation day 14 to 24 h prior to sampling at postpartum day 9. The tail suspension test (TST) and forced swim test (FST) were assessed at postpartum days 7 and 8, respectively. Unlike RU486 (mifepristone), which we have used previously ^37,38^, CORT113176 does not antagonize progesterone signaling and has no affinity for progesterone receptors ^64,79–82^.

To examine the therapeutic effects of a GR antagonist (CORT113176), an antidepressant (fluoxetine), and an allopregnanolone analog (ganaxolone) on postpartum behaviors related to mobility/despair, mice were treated with CORT113176 [80 mg/kg, *p.o.* (oral gavage with flexible polypropylene feeding tubes), dissolved in dimethyl sulfoxide and further dissolved in saline containing 0.5 % hydroxypropyl methylcellulose + 0.1 % Tween 80], fluoxetine [18 mg/kg, *p.o.* (oral gavage with flexible polypropylene feeding tubes), dissolved in deionized water], or a GABA_A_ receptor modulator ganaxolone (10 mg/kg, *i.p.*, dissolved in saline containing 0.5 % Tween 80) once daily from postpartum day 0 to 24 h prior to sampling at postpartum day 9. To avoid acute effects of the compounds immediately prior to behavioral testing, each compound was administered 24 h before each behavioral test.

Serum fluoxetine concentrations at 18 mg/kg/day in mice have been reported to be toward the upper end of the plasma concentration range seen in patients taking 20-80 mg/day of Prozac ^83^. Therefore, based on information from other publications ^66,84–88^, we chose 18 mg/kg fluoxetine for **Figure 3**. Since CORT113176 and ganoxolone have not been used clinically in human psychiatric disorders, the doses and routes chosen for **Figure 4** were determined based on those previously published by other groups ^65,67,89–91^. Since 18 mg/kg of fluoxetine and 10 mg/kg of ganoxolone did not affect behavior in **Figure 4**, the dose-response testing in **Extended Data Figure 5** systematically increased the doses of these drugs to check their efficacy. Since CORT113176 showed positive effects at 80 mg/kg in **Figure 4**, we tested both lower and higher doses in **Extended Data Figure 5**.

### Behavioral assays in mice

Two cohorts for each postpartum time point were used: the first cohort was used for TST followed by FST, and the second cohort for the three-chamber social interaction test (SIT). We conducted the FST just one day after the TST. Although exposure to the TST may interfere with the results of the FST in this experimental schedule, the effect is likely to be small. This effect would be equally present in all groups compared in this study, and this effect would not affect our conclusions. Different cohorts of mice were prepared to avoid the repeated exposure to stressful behavioral procedures (**Extended Data Figure 1**). We also prepared the third cohort of dams to perform the sucrose preference test (SPT).

#### Tail Suspension Test (TST)

Female mice were suspended using a piece of tape attached to the tail. A trimmed 1000 μl pipette tip was used over the tail to prevent biting the tape. The duration of immobility was recorded for 6 minutes at postpartum days 0, 7, and 21 and calculated as follows: 360 (sec) - struggling time (sec) = immobility time (sec).

#### Forced Swim Test (FST)

At postpartum days 1, 8, and 22 (24 h after the TST), female mice were individually placed in a transparent glass cylinder (8 cm in diameter x 20 cm high) containing water at 23 °C to a depth of 15 cm. The duration of immobility was measured for 6 minutes and calculated as follows: 360 (sec) - swimming time (sec) = immobility time (sec).

#### Sucrose Preference Test (SPT)

SPT was performed as previously described ^44,92^, with minor modifications. At postpartum days 3 or 17, regular water bottles were replaced with two 50 ml tubes (bottle “A” and bottle “B”) fitted with bottle stoppers containing two balled sipper tubes. The positions of bottles A and B were switched daily to avoid side bias, and the fluid consumed from each bottle was measured daily. During postpartum days 3-4 and 17-18, both bottles were filled with normal drinking water (W/W). During postpartum days 5-6 and 19-20, both bottles were filled with a solution of 1.5 % sucrose dissolved in drinking water (S/S). At postpartum days 7-10 or 21-24, we performed SPT. On the test days, bottle A contained 1.5 % sucrose, and bottle B contained drinking water (S/W). Sucrose preference for each mouse on each day was calculated as 100 % x [Vol A / (Vol A + Vol B)] and averaged across days for a given condition (W/W, S/S, or S/W).

#### Three-chamber Social Interaction Test (SIT)

Female mice were individually placed in the center chamber with the two adjacent chamber doors open and allowed to habituate for 10 minutes to the three chambers for three consecutive days before testing. On postpartum days 0, 7, and 21, the same mouse was placed back in the center chamber for 5 minutes with the two chamber doors closed. In the “sociability” trial of the SIT, the subject mouse encountered a novel mouse (stranger 1) in a wire cage in one chamber and an object in a wire cage in the other chamber for 10 minutes. Afterwards, the chamber doors were closed again, and the subject mouse was confined to the center chamber for five minutes. In the “social novelty recognition” trial, the subject mouse encountered stranger 1 (now familiar) and a novel mouse (stranger 2) in the previously object wire cage. The time spent sniffing each wire cage was analyzed using Ethovision computer software and manual counts. Sociability and social novelty indexes were calculated as follows: Sociability index = (Time spent sniffing a mouse cage – Time spent sniffing an object cage) / (Time spent sniffing a mouse cage + Time spent sniffing an object cage). Social novelty index = (Time spent sniffing a novel cage – Time spent sniffing a familiar cage) / (Time spent sniffing a novel cage + Time spent sniffing a familiar cage) ^93^. Of note, the data presented in **Extended Data Figure 2** of the present study, where the experiments were carried out within a single day, differed from the data obtained in a separated study, where the experiments were conducted over two consecutive days ^41^.

### Quantitative reverse transcription PCR

The PVN and pituitary were rapidly dissected out 24 h following the last behavioral test. Total RNA was isolated by using the RNeasy Mini Kit and converted into cDNA with the SuperScript^TM^ III First-Strand System for RT-PCR Kit. Real-time PCR reactions were carried out in duplicate using the 1 × TaqMan master mix, 1 × TaqMan probes for each gene (Crh, Crhr1, Gr, and β-actin), and 200 ng of cDNA template in a total volume of 20 μL. The β-actin TaqMan probe was used as the internal control. Real-time PCR was performed on a Bio-Rad CFX96 Touch Real-Time PCR Detection System with standard PCR conditions (50°C for 2 min, 95°C for 10 min, 95°C for 15 s, and 60°C for 1 min for 40 cycles). The expression levels were calculated as described previously ^94^.

### Measurement of plasma hormone levels in mice

Blood was collected from the inferior vena cava under isoflurane anesthesia between 9 am and 11 am, 24 h following the FST. Levels of plasma corticosterone, estradiol, progesterone, oxytocin, and prolactin were assessed using commercially available enzyme-linked immunosorbent assay (ELISA) kits ^37^ and measured at four time points [virgin, late pregnancy (gestation day 19), 0 week postpartum (postpartum day 2), and 1 week postpartum (postpartum day 9)]. Plasma corticosterone was also measured at 3 weeks postpartum (postpartum day 23) following the last behavioral testing. Plasma samples from different cohorts of mice at each time point were prepared to avoid the repeated exposure to stressful behavioral procedures. To measure the function of the HPA axis negative feedback loop, a dexamethasone (DEX) suppression test was performed at postpartum day 8. Dexamethasone is a corticosteroid that, at low doses, acts on the anterior pituitary gland to suppress adrenocorticotropic hormone (ACTH) release, which inhibits corticosterone release from the adrenal gland ^54–56^. DEX at low doses cannot pass the blood-brain barrier and can provide insight into the functionality of the HPA axis ^52,53^. DEX (0.1 mg/kg, *i.p*) was administered to mice six hours prior to collecting blood samples at 5 pm on postpartum day 8 ^95,96^. Plasma ACTH and corticosterone levels after treatment with DEX were assessed using commercially available EIA kits as published ^37^.

#### Blood glucose testing and measurement of weight of thymus, spleen, and visceral fat in mice

Mice were deprived of food beginning at 8 am for 6 hours at postpartum day 8. After a 6-hr fasting, basal levels of blood glucose were measured using an Accu-Chek Softclix glucometer ^97,98^. Then glucose (2.0 g/kg, *i.p.*) was administered to mice. Blood glucose levels 30, 60, 90, and 120 min after the glucose injections were measured using the glucometer ^97,98^. We collected blood from tail vein. After the blood glucose testing, thymus, spleen, and visceral fat were weighted.

### Human subjects

Study participants were recruited through the Johns Hopkins Women’s Mood Disorders Center and written informed consent was obtained by all patients. 116 pregnant women (18 years or older) with a history of mental illnesses underwent psychological interviews with the Structured Clinical Interview for DSM-IV Axis 1 Disorders (SCID) by study psychiatrists, and blood draws were conducted between 8:45 am and 4:45 pm at four time points across pregnancy and the postpartum period (2^nd^ and 3^rd^ trimester, 2 weeks postpartum, and 6 weeks postpartum). PPD was defined as developing major depression within six weeks of delivery and not during pregnancy ^99^. “History” refers to their psychiatric history up until their first visit (**Extended Data Table 1**). Subjects were also assessed regarding childhood home environment, childhood behavior, and traumatic events (occurring before 17 years of age) (**Extended Data Table 1**). Abnormal childhood home environment included physical abuse of kids or parents, sexual abuse, lots of fighting, food insecurity, or other stressful family environment. Abnormal childhood behavior included truancy, running away, setting fires, getting kicked out of school, harming of animals, stealing, or other antisocial types of behaviors. Traumatic events included death of a close friend/family member, major upheaval between parents, sexual assault, non-sexual assault, extreme illness/injury, or other major life events. Each traumatic event was scored from 1-7 based on the individual’s subjective rating of how traumatic the event was. A score of zero indicated no event occurred for that participant.

### Measurement of plasma cortisol levels in humans

Plasma cortisol levels were assessed using a commercially available ELISA kit. Confounding effects of age, race, blood draw time, and medication (anti-psychotic, antidepressant SSRIs, and non-SSRI antidepressants) were adjusted for.

### Statistical Analysis

Statistical analyses were performed using commercial software (GraphPad Prism 7 and 9, GraphPad Software, Inc., IBM SPSS statistics 26, IBM, and R version 3.5.1) (**Extended Data Table 2**). Normality of data sets were tested using Shapiro-Wilk’s test. *For* normally distributed data, statistical differences between two groups were calculated using the Student’s t-test. Statistical differences among three groups or more were determined using a two-way analysis of variance (ANOVA) and three-way ANOVA with repeated measures, followed by a Bonferroni post hoc test. Corrections for multiple comparisons were made when appropriate. For non-normally distributed data, statistical differences between two groups were calculated using the Mann-Whitney test. Statistical differences among three groups or more were determined using a Kruskal-Wallis test followed by a Bonferroni post hoc test. Spearman’s correlation or Pearson’s correlation were used to examine correlations between plasma corticosterone levels and behavioral changes in TST and FST. For human data, the Fisher exact test was used to check if two categorical variables are related, for example, if PPD is related to traumatic events. Linear regression was employed to evaluate the contributions of risk factors to PPD. The t-values from the liner regression were used to estimate the impact of these factors on PPD. In addition, linear regression was conducted to evaluate the associations between cortisol values and PPD, with age, race, medication, and blood draw time as covariates. A value of *p* < 0.05 was considered statistically significant. All data are expressed as the mean ± SEM.

## Acknowledgements

We thank the Corcept Therapeutics Inc., especially Dr. Hazel Hunt and Dr. Joseph K. Belanoff for providing the specific GR antagonist CORT113176 and technical support for its use in an academic study. We also thank clinical study participants and staff members. We appreciate Ms. Courtney Erdly and Ms. Dania Mallah for their technical help, and Dr. Melissa A. Landek-Salgado for scientific reading. This work was supported by the National Institute of Health [MH-092443 (A.S.), MH-094268 Silvio O. Conte center (A.S.), K99MH-094408 (M.N.), MH-105660 (A.S.), MH-107730 (A.S.), DA-040127 (A.S. and M.N.), and MH-116869 (M.N.)], NARSAD (A.S. and M.N.), Stanley (A.S.), S-R/RUSK (A.S.), JST PRESTO JPMJPR14M6 (M.N.), and UAB Psychiatry startup funds (M.N.).

## Author contributions

**M.N.** conceived and designed the project, and supervised **S.L.**, **D.J.W.**, **J. F-O.**, **K.K.**, and **A. A.** for practical experiments with the help of **S.K.** and the guidance of **A.S.**. Experiments were performed by **M.N.**, **S.L.**, **D.J.W.**, **J. F-O.**, **K.K.**, and **A.A.**. Data was analyzed by **M.N.**, **K.Y.**, **J. F-O.**, and **K.K.**. The first manuscript draft was written by **M.N.** and **A.S.** with input from all the authors. **M.N.** and **A.S.** revised and edited the manuscript for the final version. **G.S.W.** provided expertise of endocrinology in a translationally relevant manner. **J.L.P.** provided human plasma samples with participants’ information as well as expertise of postpartum mood disorders.

## Competing interests

The authors declare no competing interests.

## Materials and Correspondence

Correspondance and requests for materials should be addressed to Akira Sawa.

---

**Extended Data Figure 1.**
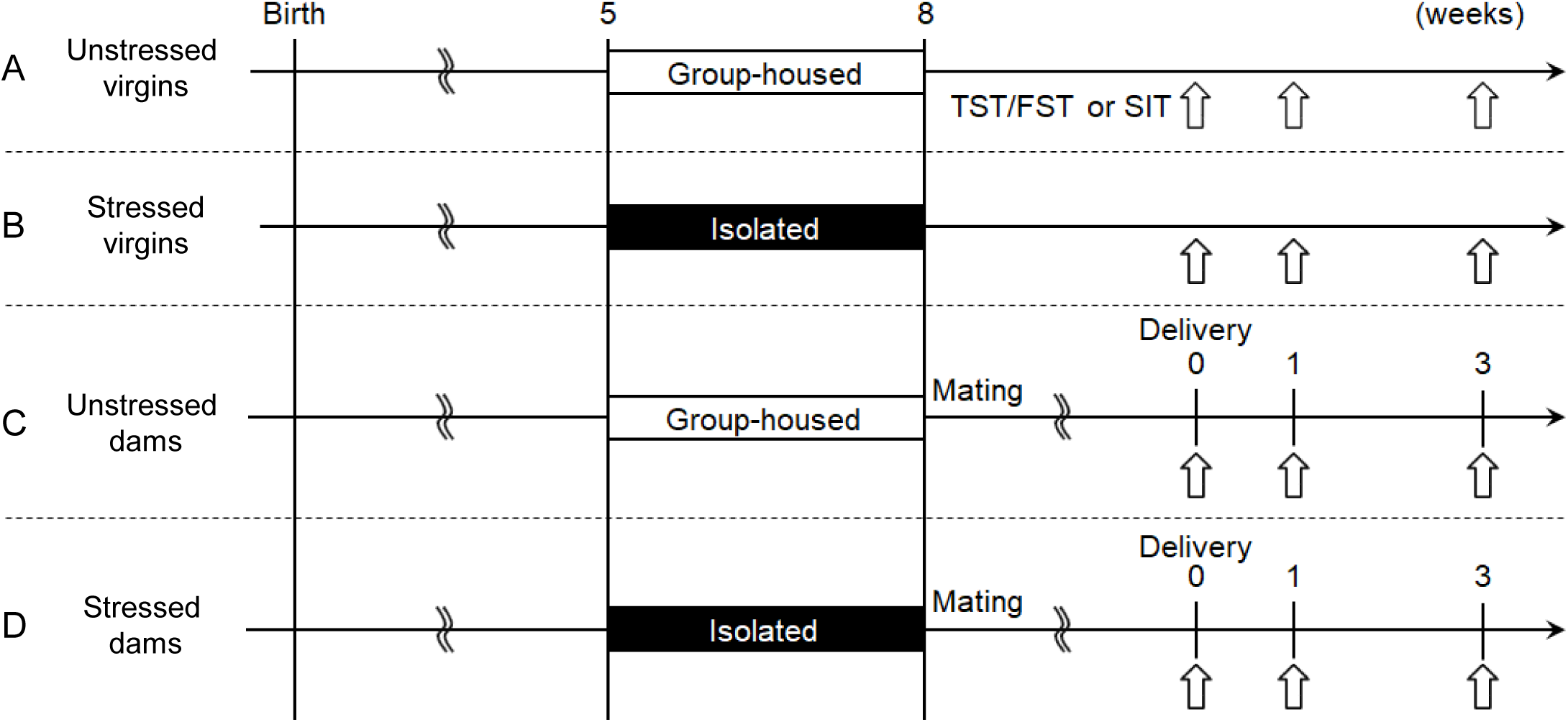
Experimental schedule of the preclinical study. **A**, Virgin female mice were group-housed. **B**, Virgin female mice were isolated from 5 to 8 weeks of age. **C**, Virgin female mice were group-housed, mated with a male mouse, and gave birth to pups. **D**, Virgin female mice were isolated from 5 to 8 weeks of age, mated with a male mouse, and gave birth to pups. TST, tail suspension test; FST, forced swim test; SIT, three-chamber social interaction test. Different cohorts of mice subjected to behavioral tests at 0 week, 1 week, and 3 weeks postpartum were studied to avoid the repeated exposure to stressful behavioral procedures.

**Extended Data Figure 2.**
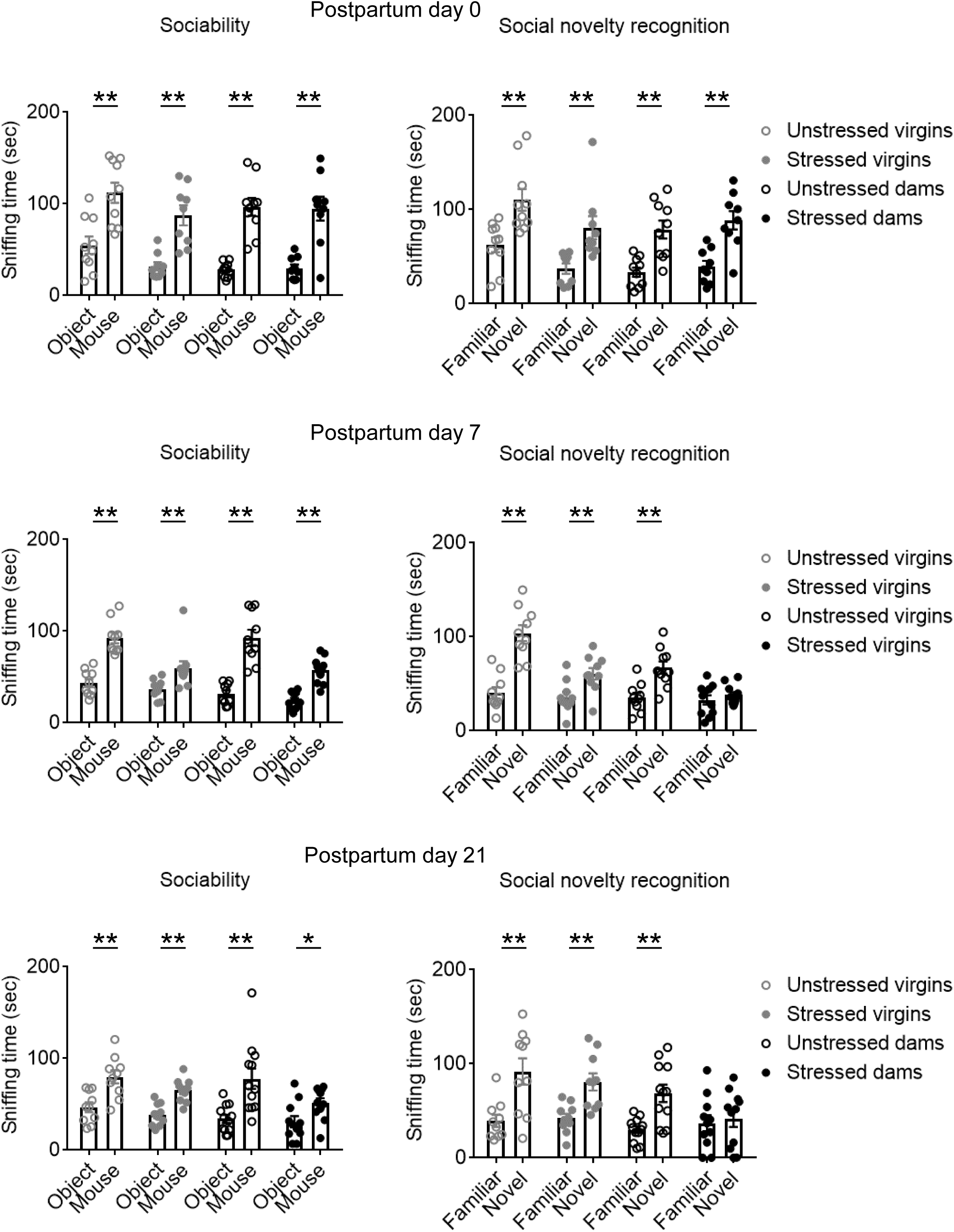
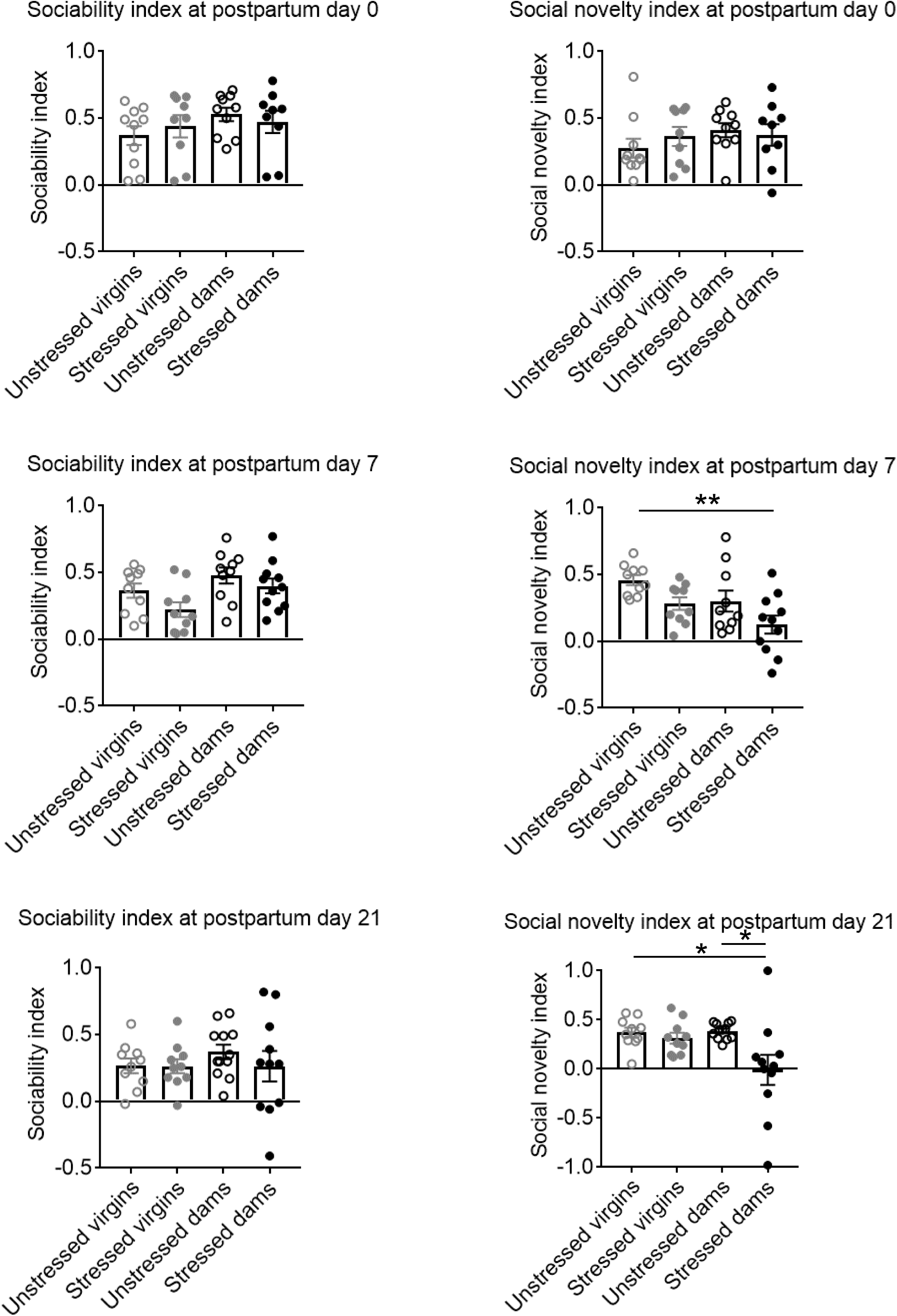
Long-lasting behavioral changes in the social interaction test in dams exposed to adolescent social isolation. Sniffing time and indexes of sociability and social novelty recognition during the three-chamber social interaction test were measured at postpartum days 0, 7, and 21. No changes in sociability or social novelty recognition among the four groups were observed at postpartum day 0. Behavioral changes in social novelty recognition, but not sociability, were observed in stressed dams at postpartum days 7 and 21. N=9-11. Values are represented as mean ± SEM; \*\**P*<0.01 and \**P*<0.05.

**Extended Data Figure 3.**
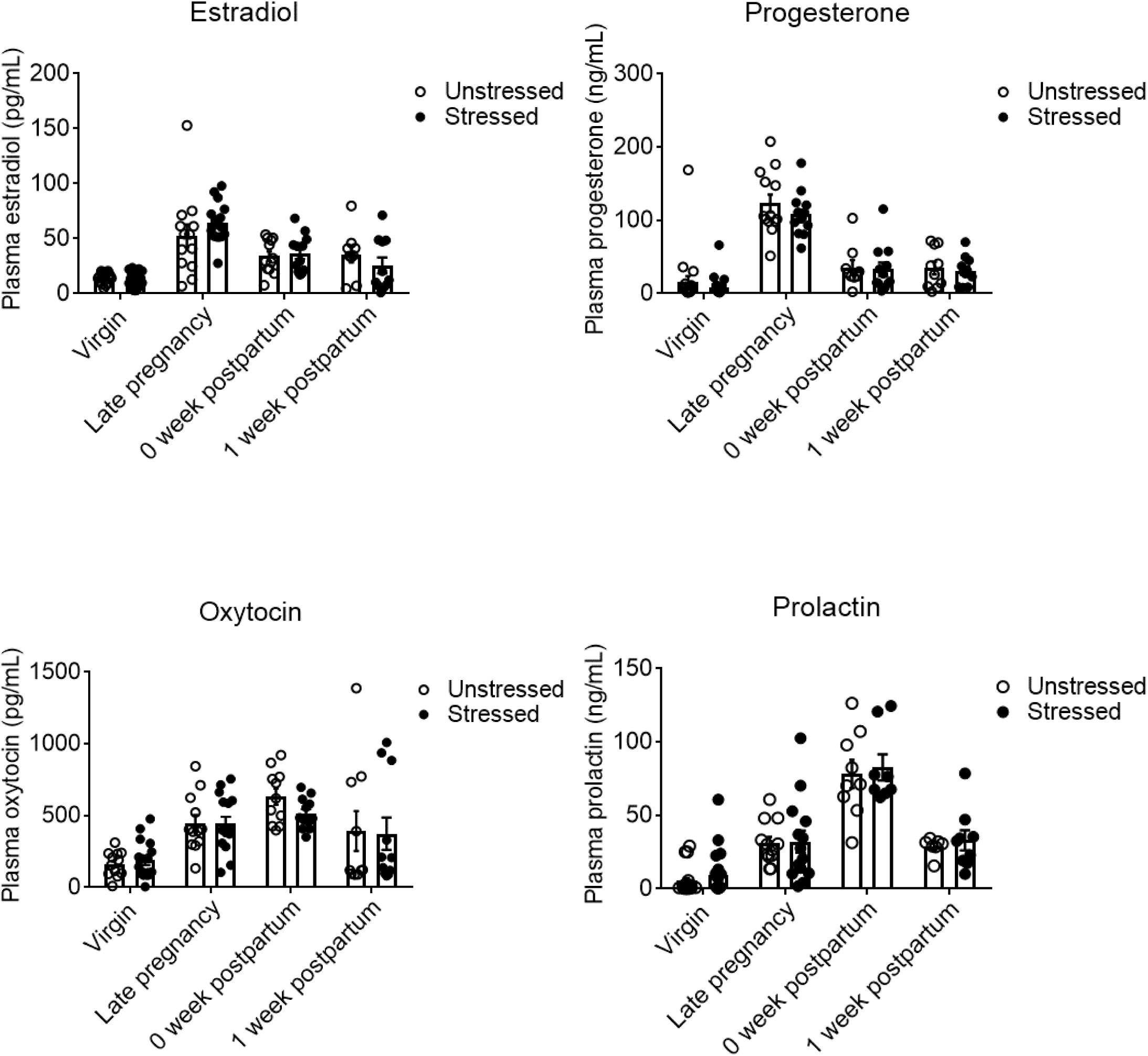
No change in the levels of plasma estradiol, progesterone, oxytocin, and prolactin in mice exposed to adolescent social isolation. Levels of estradiol, progesterone, oxytocin, and prolactin in plasma were measured at four-time points (virgin, late pregnancy, 0 week postpartum, and 1 week postpartum). No differences in the levels of estradiol, progesterone, oxytocin, and prolactin were observed between unstressed and stressed mice at any time point. N=9-20. Values are represented as mean ± SEM.

**Extended Data Figure 4.**
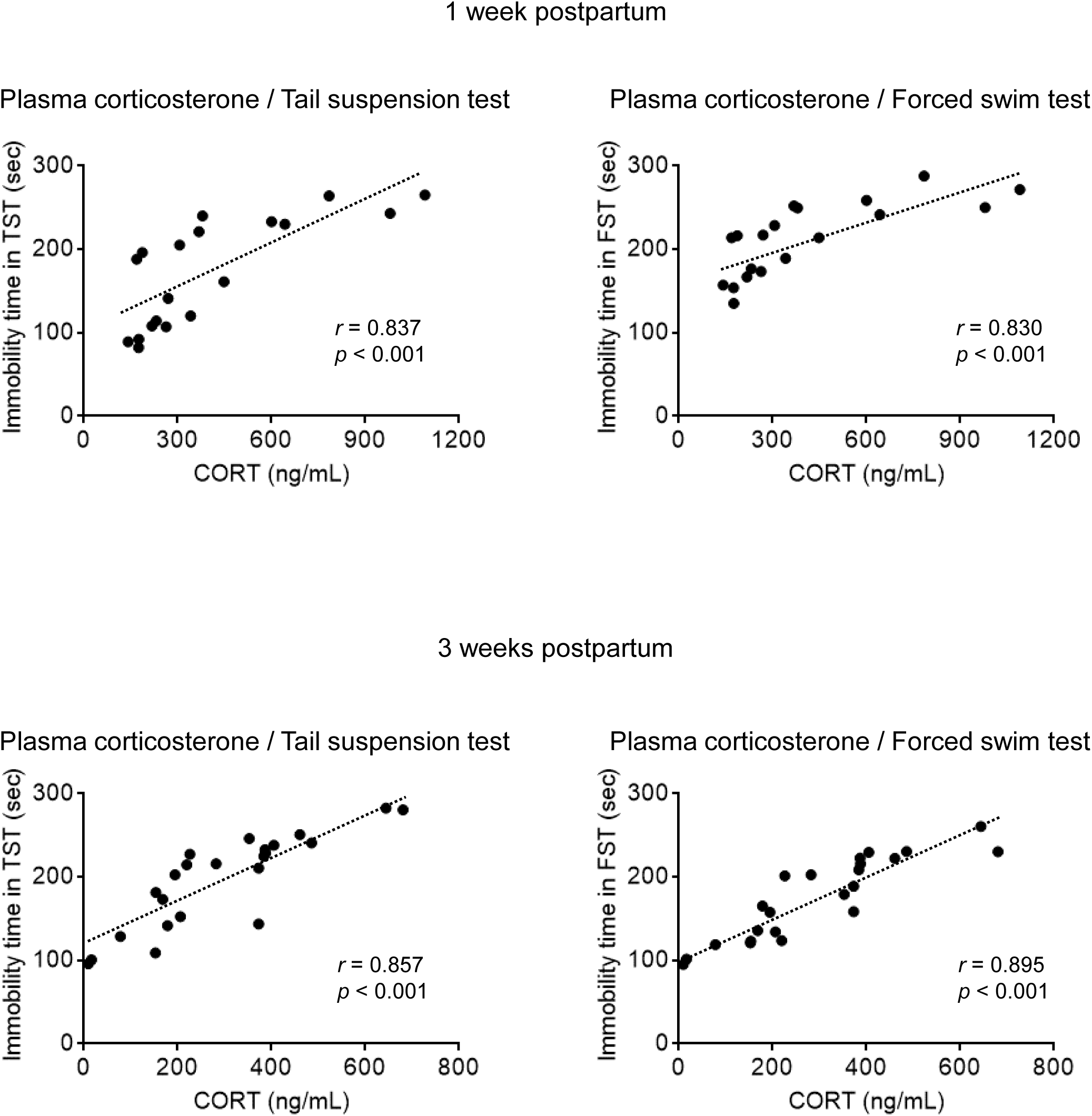
Positive correlation between plasma corticosterone levels and immobility time in the tail suspension and forced swim tests at 1 week and 3 weeks after delivery. The levels of plasma corticosterone (CORT) were positively correlated with immobility time in the tail suspension test (TST) and forced swim test (FST) at 1 week and 3 weeks after delivery. N=19-23. Spearman and Pearson rank correlation coefficients were examined for the data at 1 week and 3 week postpartum, respectively.

**Extended Data Figure 5.**
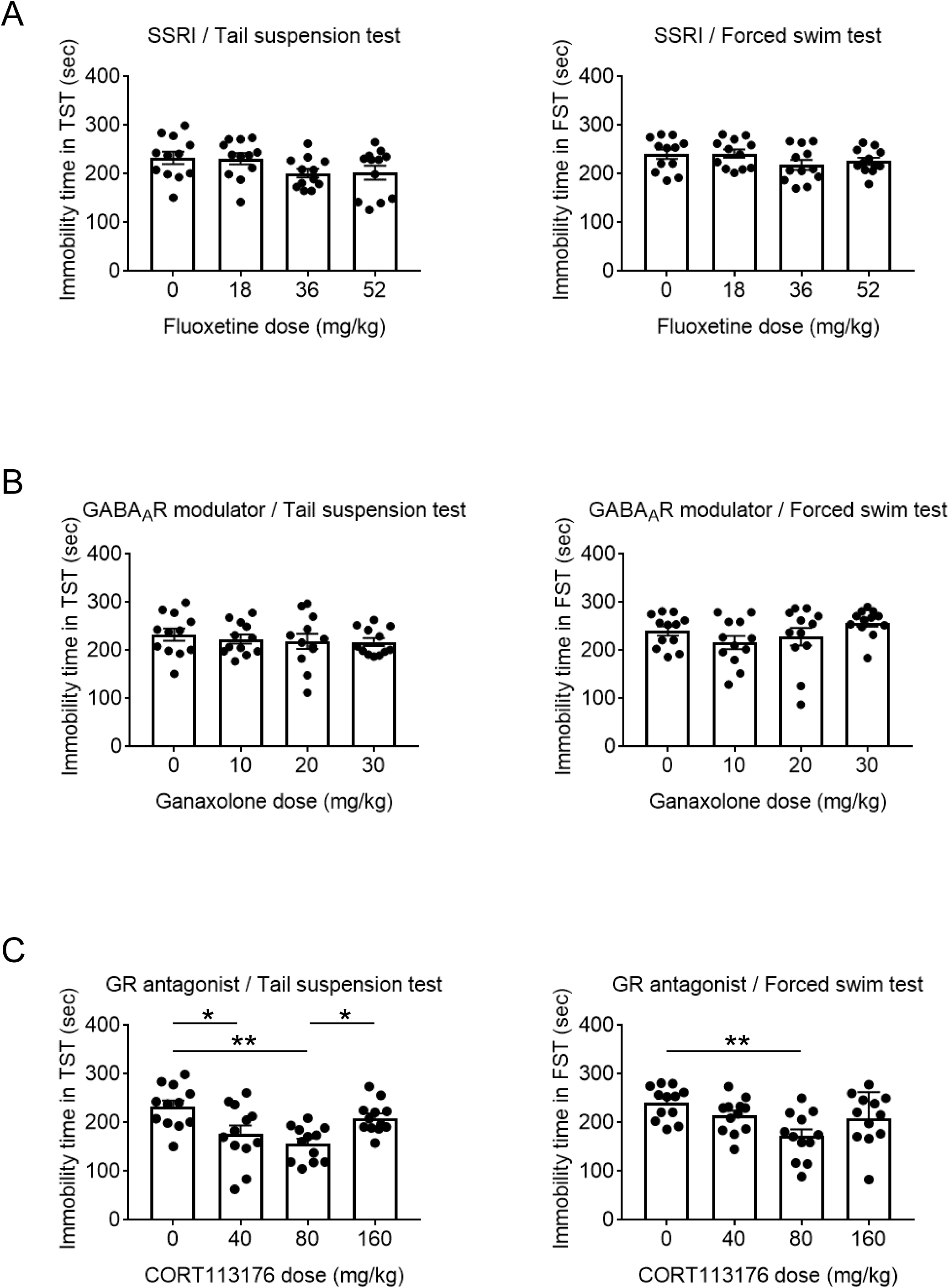
Dose-response effects of a SSRI, a GABA_A_ receptor modulator, or a GR antagonist on behavioral changes in dams exposed to adolescent social isolation. Stressed dams were treated with a SSRI fluoxetine (18, 36, 52 mg/kg, *p.o.*), a GABA_A_ receptor modulator ganaxolone (10, 20, 30 mg/kg, *i.p.*), or a GR antagonist CORT113176 (40, 80, 160 mg/kg, *p.o.*) once daily from postpartum day 0 to 24 h prior to sampling at postpartum day 9. Post-delivery treatment with only the GR antagonist at 40 and 80 mg/kg ameliorated the behavioral changes in the tail suspension test on postpartum day 7 in stressed dams. Post-delivery treatment with only the GR antagonist at 80 mg/kg ameliorated the behavioral changes in forced swim test on postpartum day 8 in stressed dams N=12. Values are represented as mean ± SEM; \*\**P*<0.01 and \**P*<0.05.

**Extended Data Figure 6.**
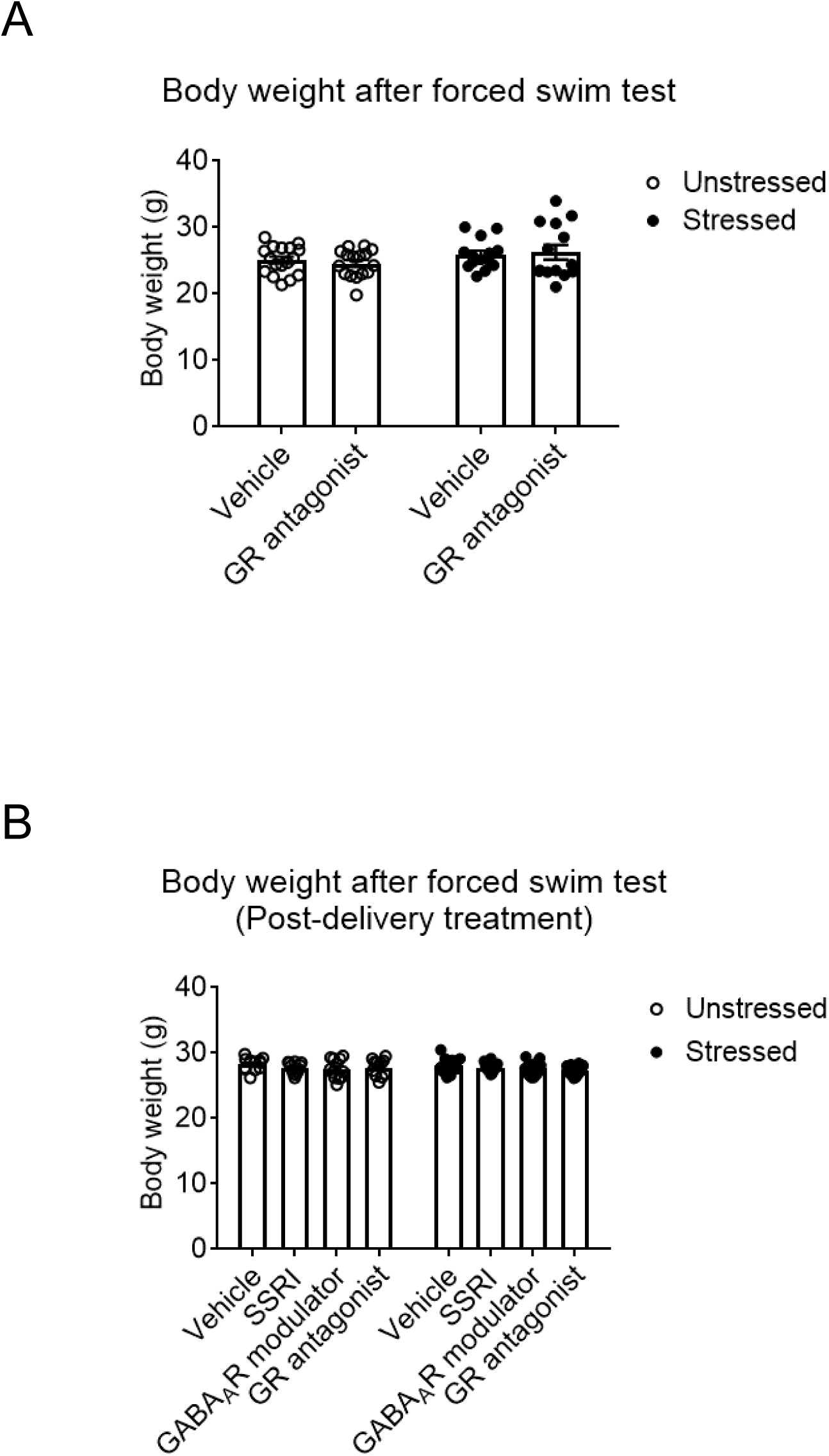
No effect of a GR antagonist, a SSRI, and an allopregnanolone analog on body weight. **A**, CORT113176 (80 mg/kg, *p.o.*, once daily from gestation day 14 to 24 h prior to behavioral testing), a selective GR antagonist, did not affect body weight after the forced swim test at postpartum day 8 in either group. **B**, Post-delivery treatment with a SSRI fluoxetine (18 mg/kg, *p.o.*), a GABA_A_ receptor modulator ganaxolone (10 mg/kg, *i.p.*), or CORT113176 (80 mg/kg, *p.o.*) once daily from postpartum day 0 to 24 h prior to behavioral testing did not affect body weight after the forced swim test at postpartum day 8 in any of the groups. N=12. Values are represented as mean ± SEM.

**Extended Data Figure 7.**
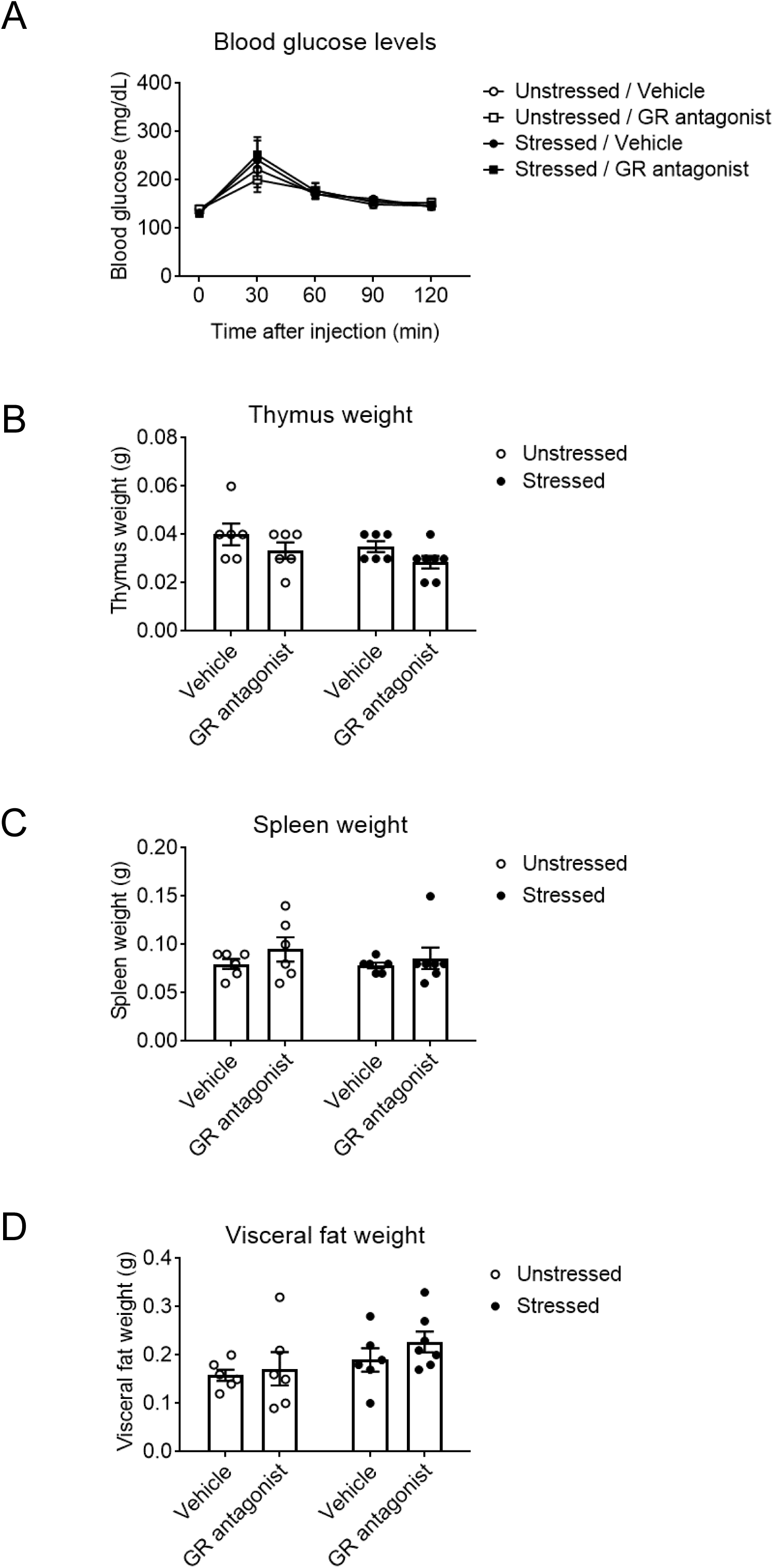
No effect of a GR antagonist on blood glucose levels and weight of the thymus, spleen, and visceral fat. **A**, After fasting for 6 hours, glucose (2 g/kg) was administered intraperitoneally and blood glucose levels at 0, 30, 60, 90, and 120 minutes were examined on postnatal day 8. CORT113176 (80 mg/kg, *p.o.*, once daily from postpartum day 0 to 24 h prior to the blood glucose testing at postpartum day 8), a selective GR antagonist, did not affect blood glucose levels in either group. **B-D**, Thymus, spleen, and visceral fat weights were measured after blood glucose testing at postpartum day 8. CORT113176 (80 mg/kg, *p.o.*, once daily from postpartum day 0 to 24 h prior to the blood glucose testing at postpartum day 8) had no effect on weight of the thymus (B), spleen (C), and visceral fat (D) in any groups. N=6-7. Values are represented as mean ± SEM.

**Extended Data Table 1.** Human study participants’ characteristics.

| Study participants (N=116) |  |  |
| --- | --- | --- |
| **Characteristics** | **N** | **%** |
| *Age, years* |  |  |
| <24 | 1 | 0.9 |
| 25-29 | 23 | 19.8 |
| 30-34 | 59 | 50.9 |
| 35-39 | 26 | 22.4 |
| >40 | 7 | 6.0 |
| *Ethnicity* |  |  |
| Hispanic or Latino | 5 | 4.3 |
| Not Hispanic or Latino | 111 | 95.7 |
| *Race* |  |  |
| Black or African-American | 11 | 9.5 |
| White | 98 | 84.5 |
| Asian or Pacific Islander | 5 | 4.3 |
| **Personal history of psychiatric disorders** | **N** | **%** |
| Major Depression | 61 | 52.6 |
| General Anxiety | 29 | 25.0 |
| Substance Abuse | 9 | 7.8 |
| Other | 42 | 36.2 |
| No history | 34 | 29.3 |
| **Psychiatric medication use** | **N** | **%** |
| Anti-psychotic | 3 | 2.6 |
| SSRI | 21 | 18.1 |
| Non-SSRI antidepressant | 6 | 5.2 |
| **Home environment** | **N** | **%** |
| Normal | 83 | 71.6 |
| Other | 33 | 28.4 |
| **Childhood behavior** | **N** | **%** |
| Normal | 96 | 82.8 |
| Other | 20 | 17.2 |
| **Traumatic event(s) before 17 years old** | **# of occurrences** | **%** |
| Death of a close friend/family member | 44 | 37.9 |
| Major upheaval between parents | 30 | 25.9 |
| Sexual assault | 18 | 15.5 |
| Non-sexual assault | 13 | 11.2 |
| Extremely ill/injured | 9 | 7.8 |
| Major upheaval that shaped life/ personality | 34 | 29.3 |

**Extended Data Table 2.**
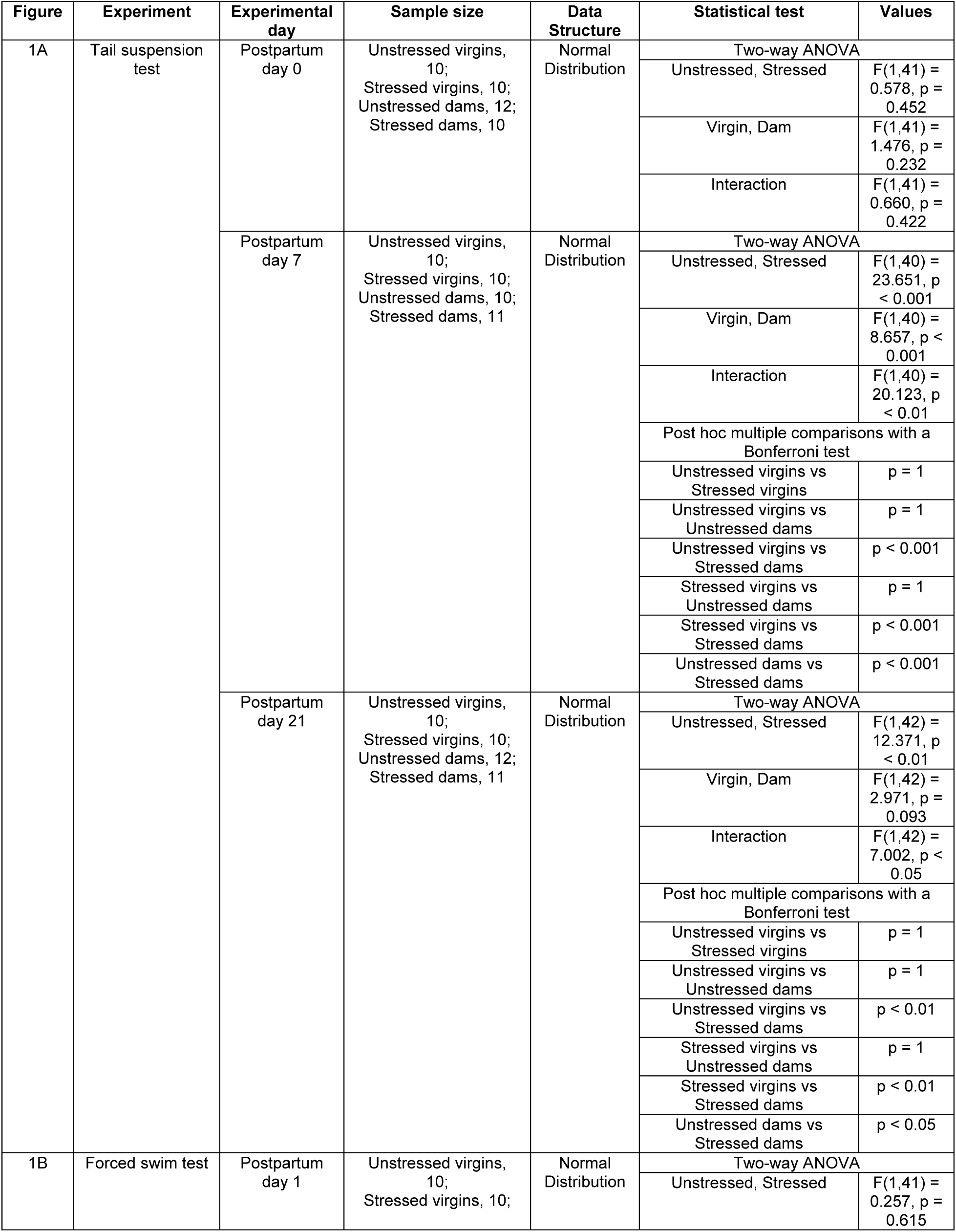
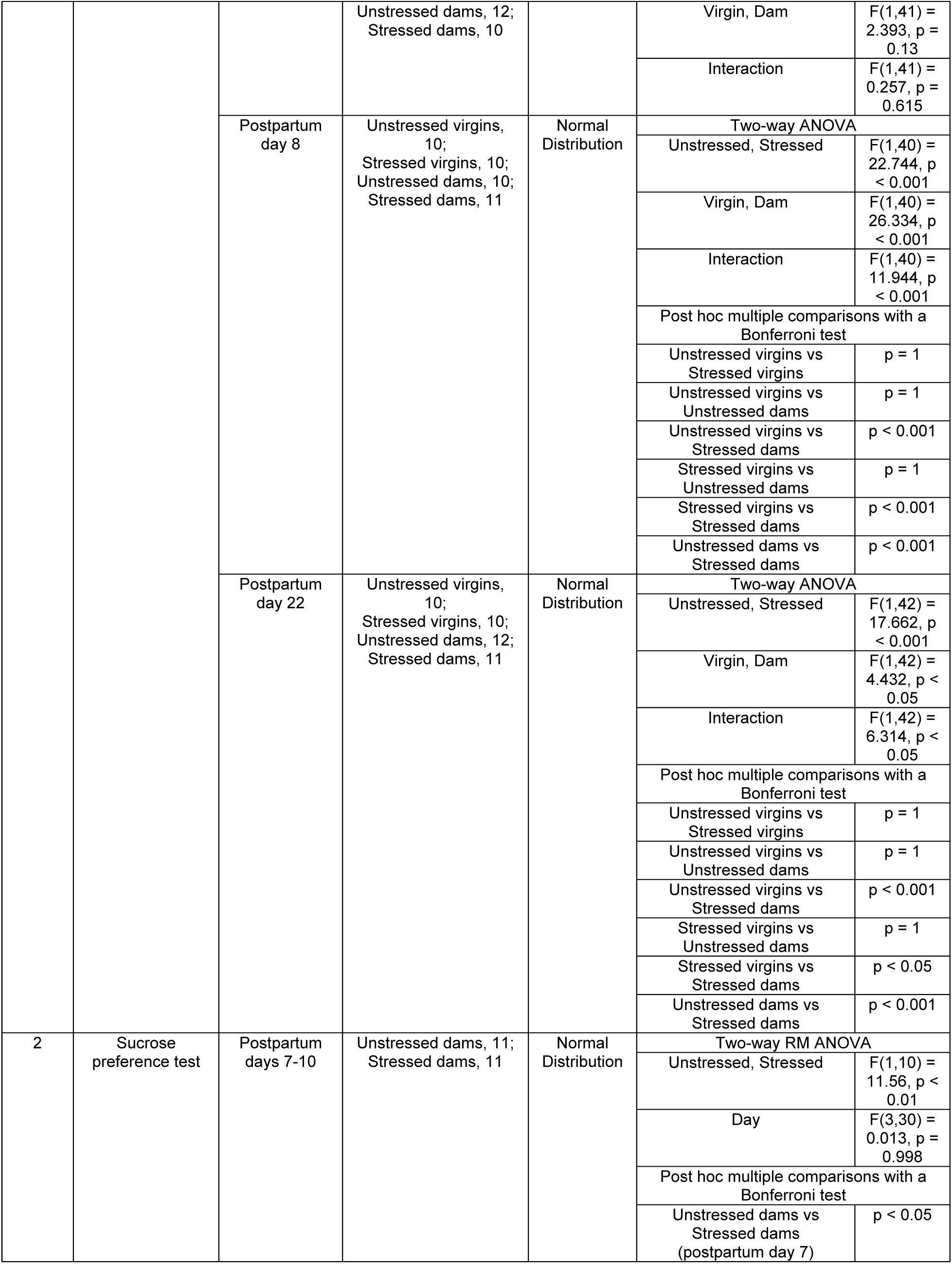
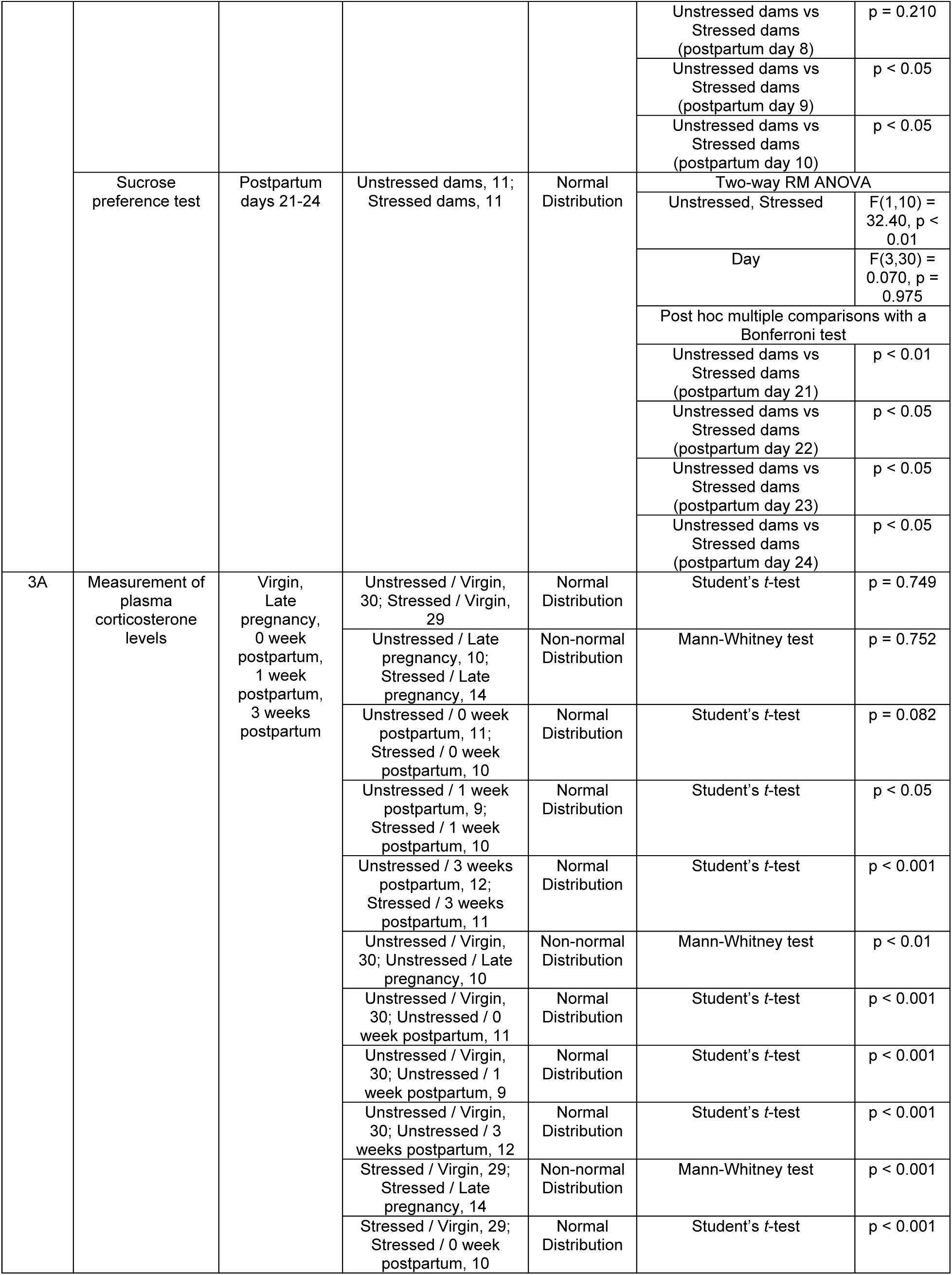
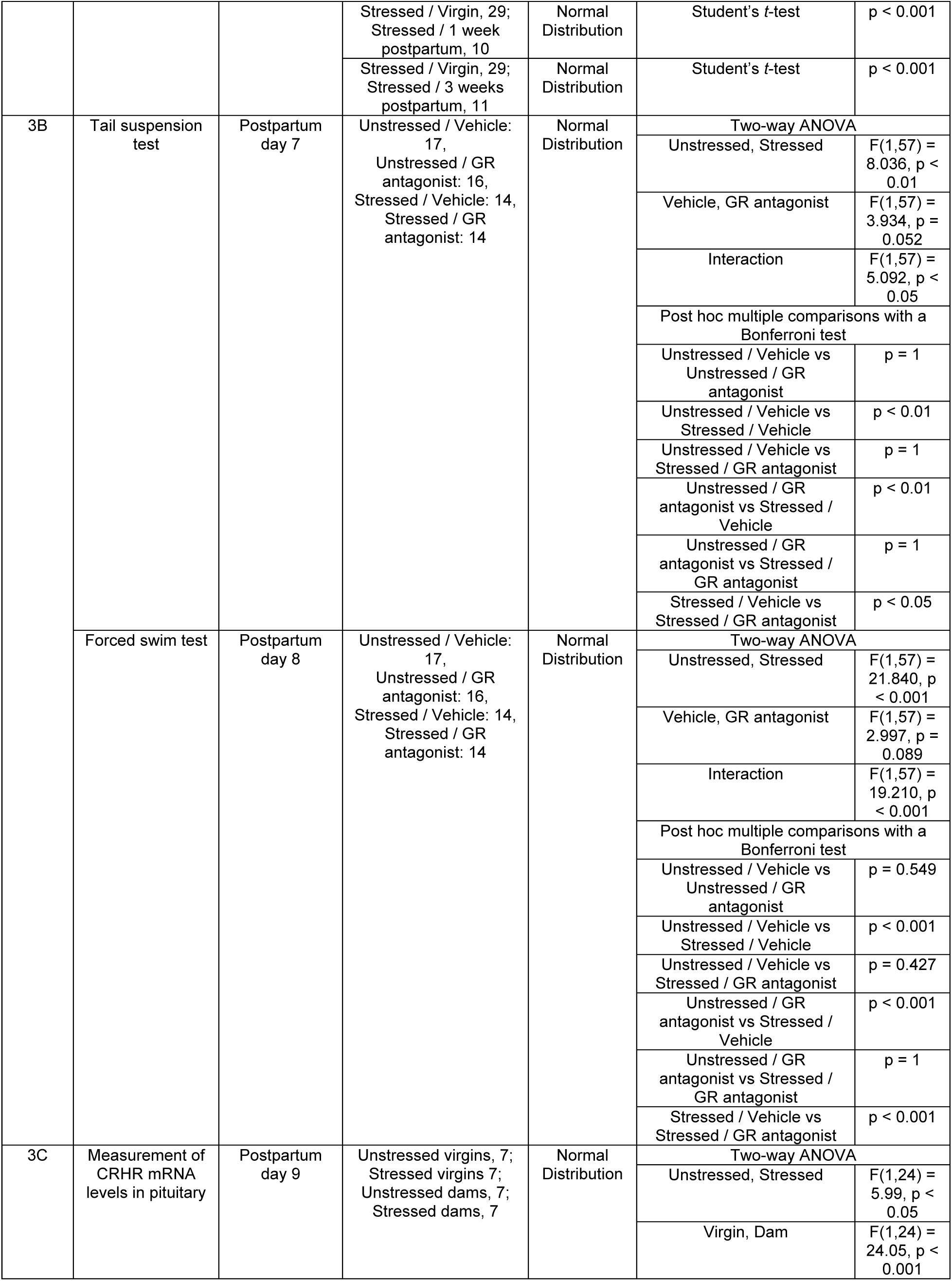
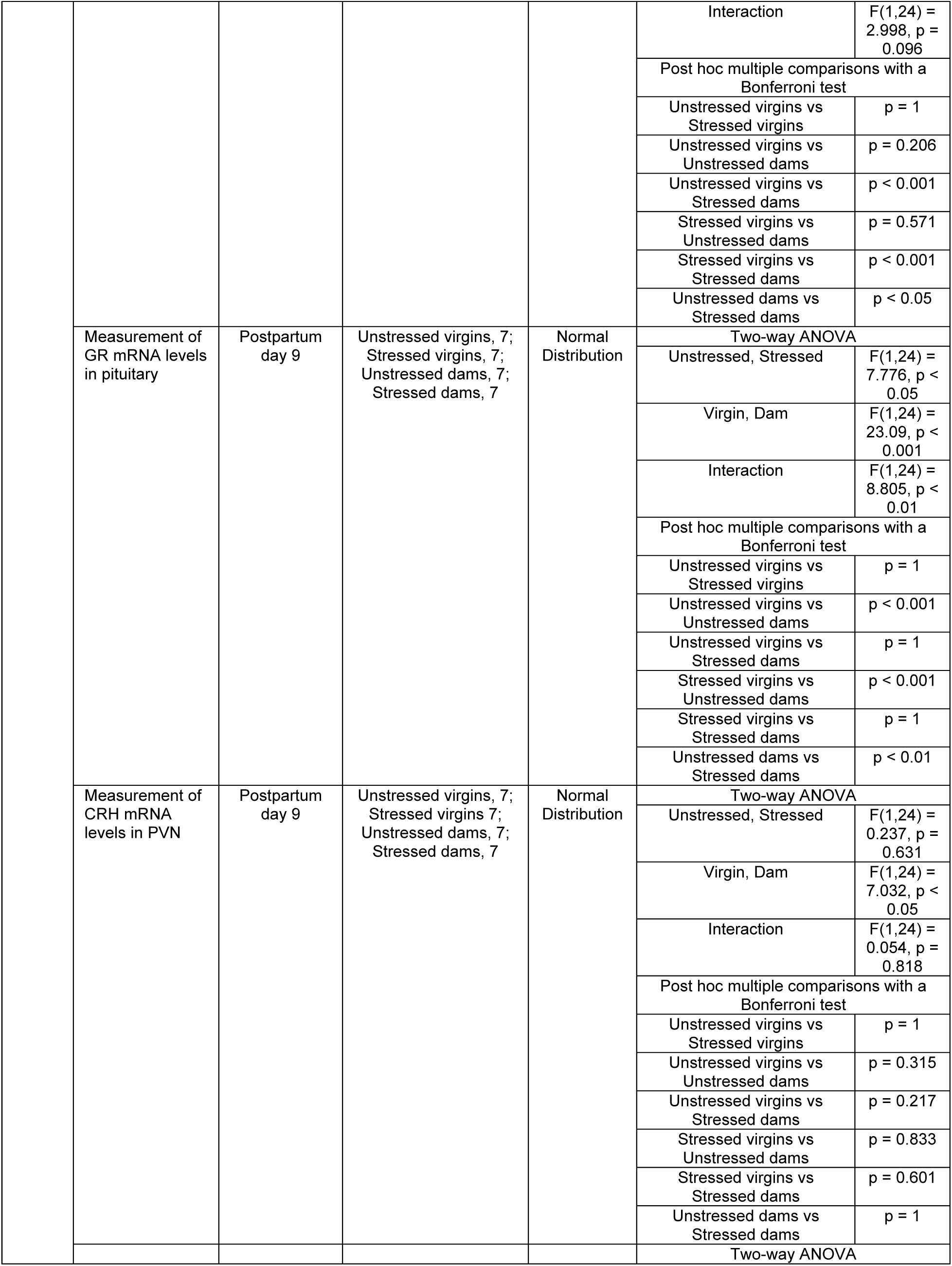
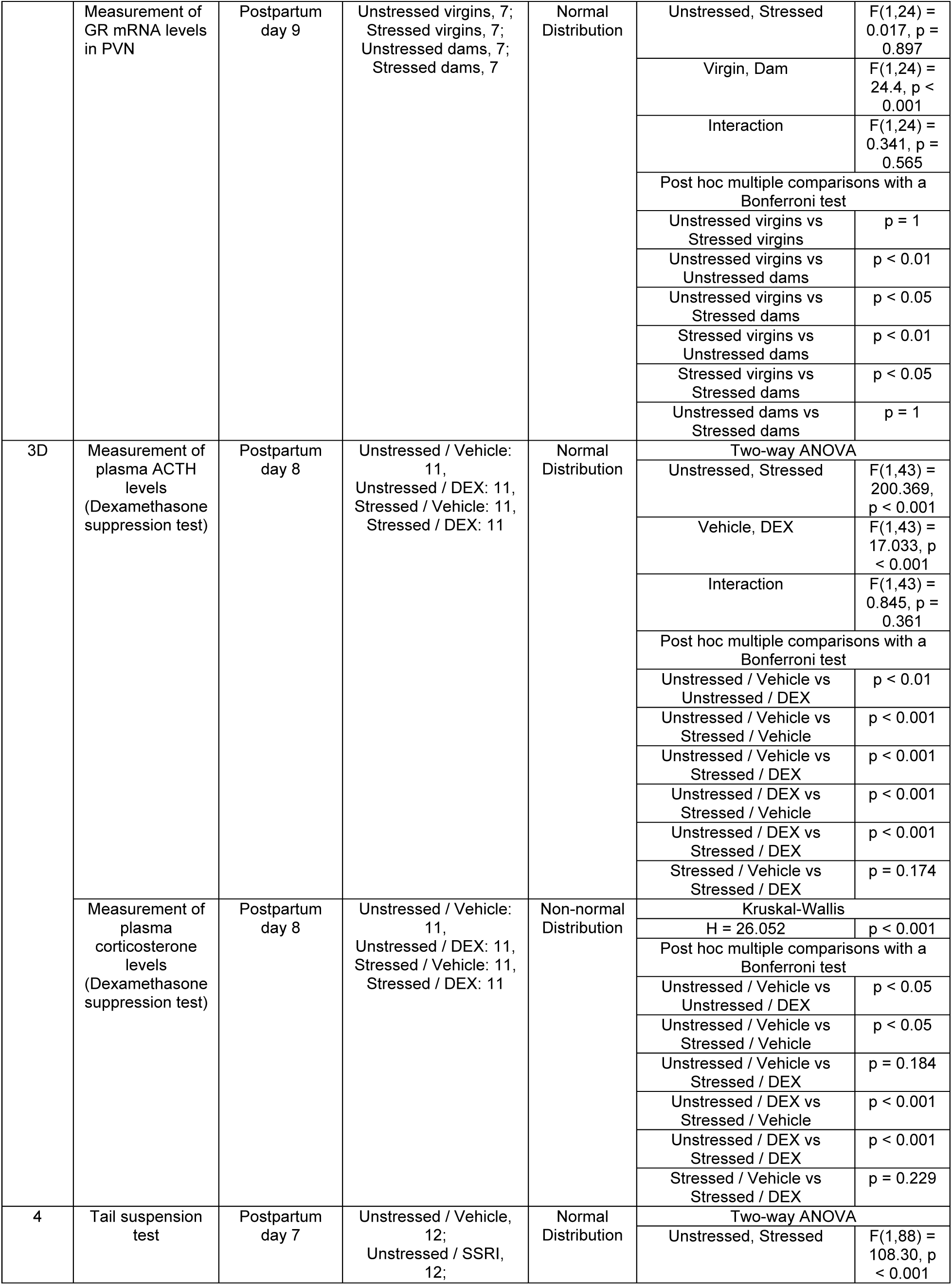
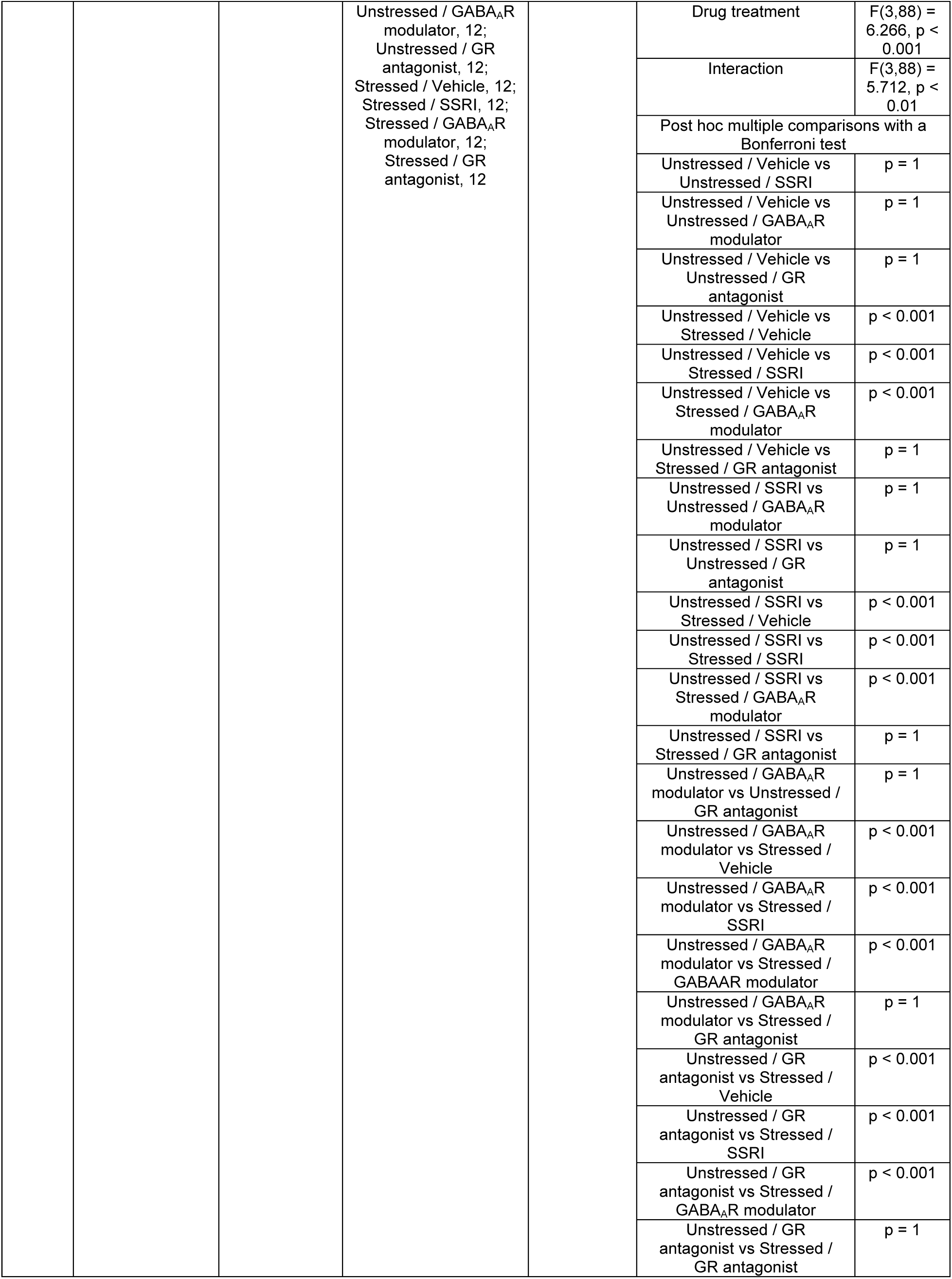
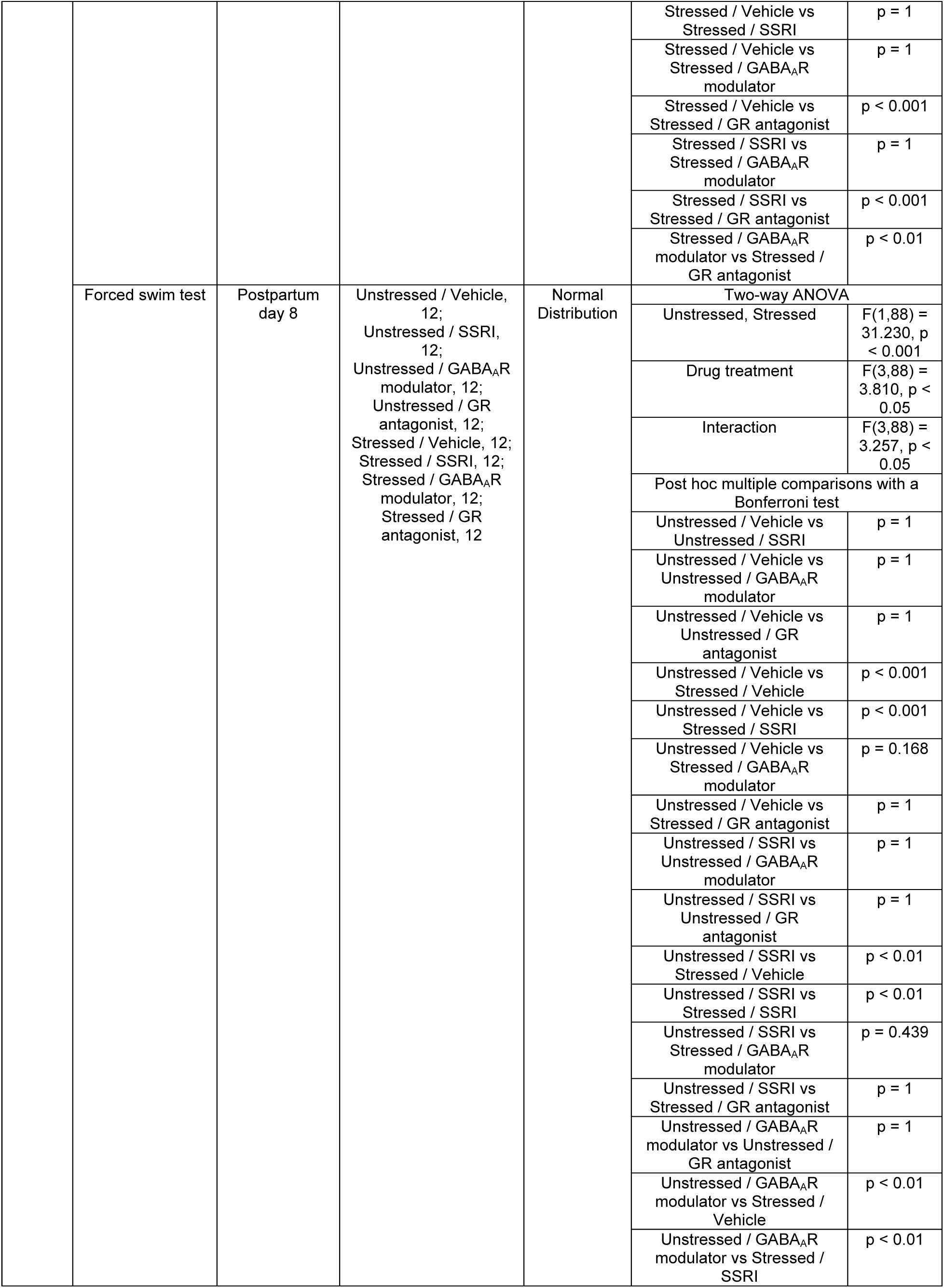
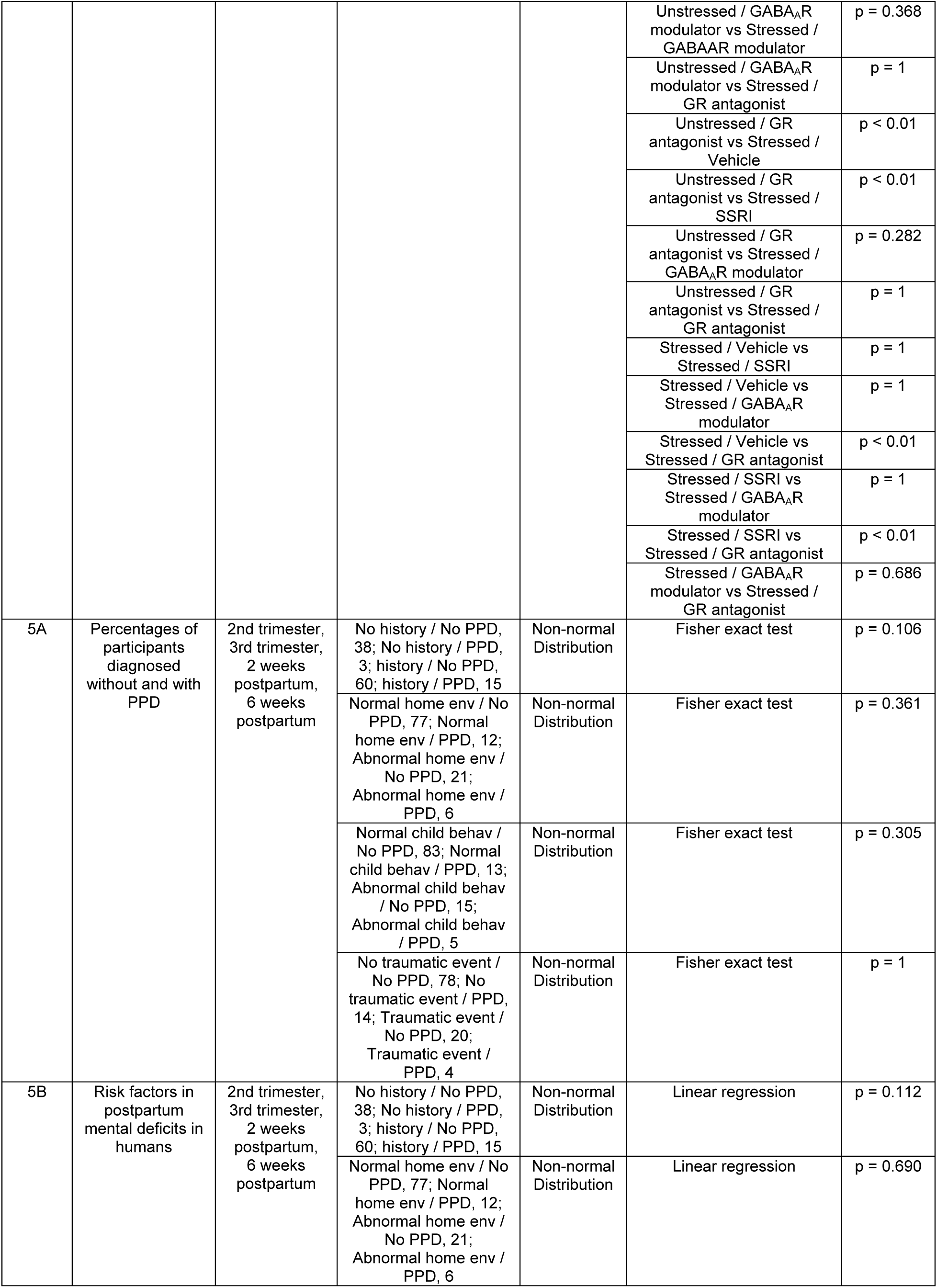
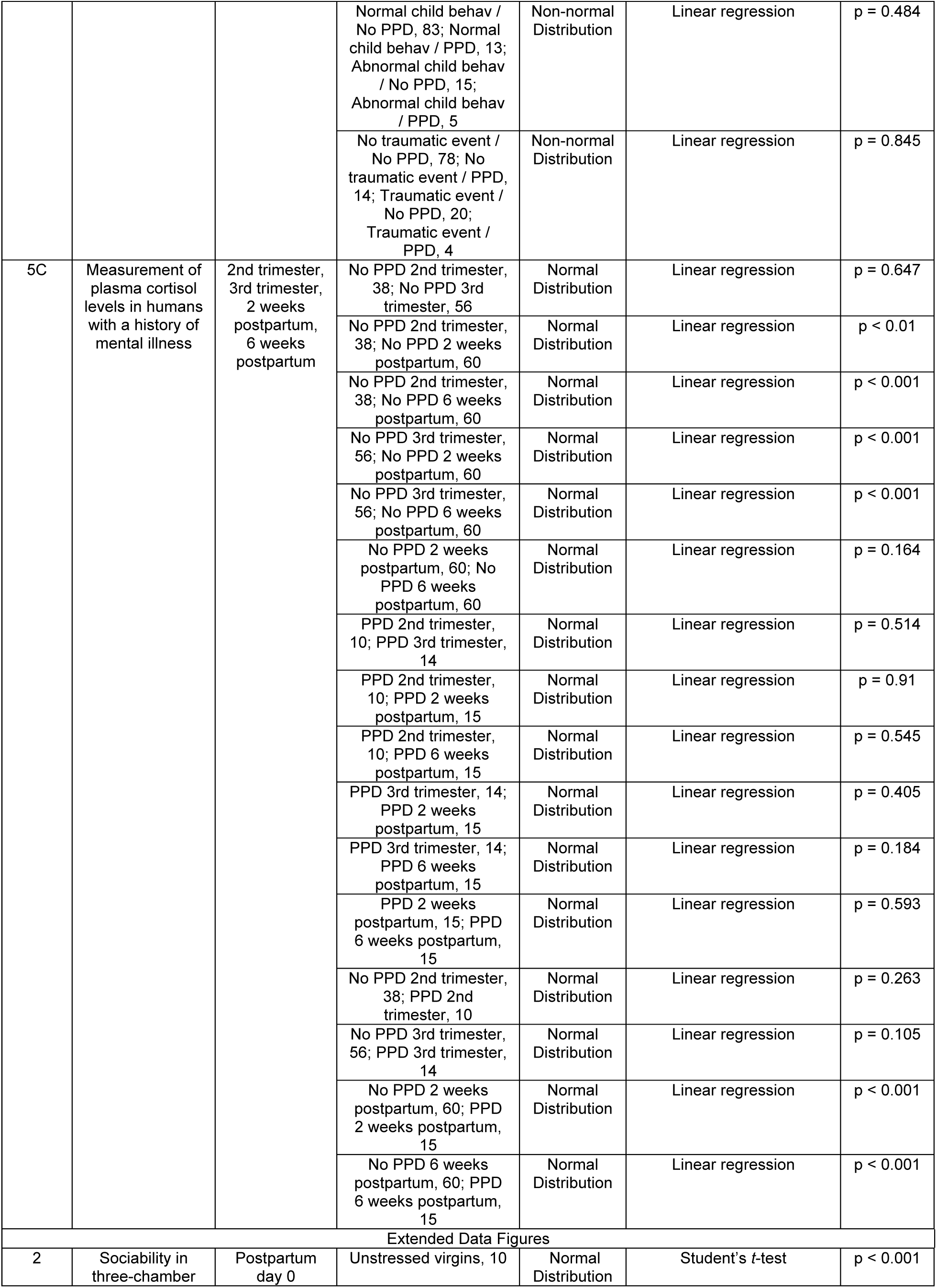
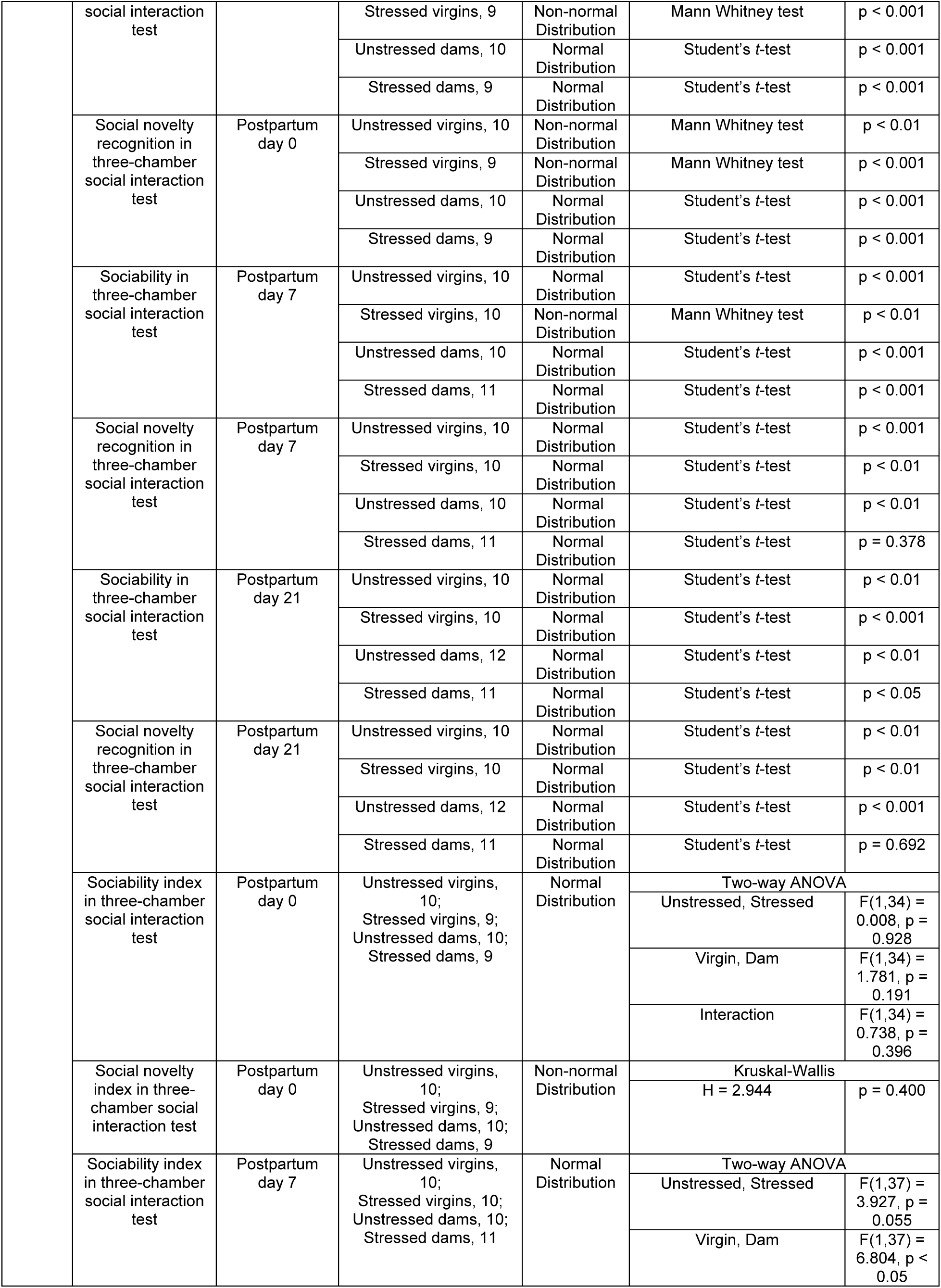
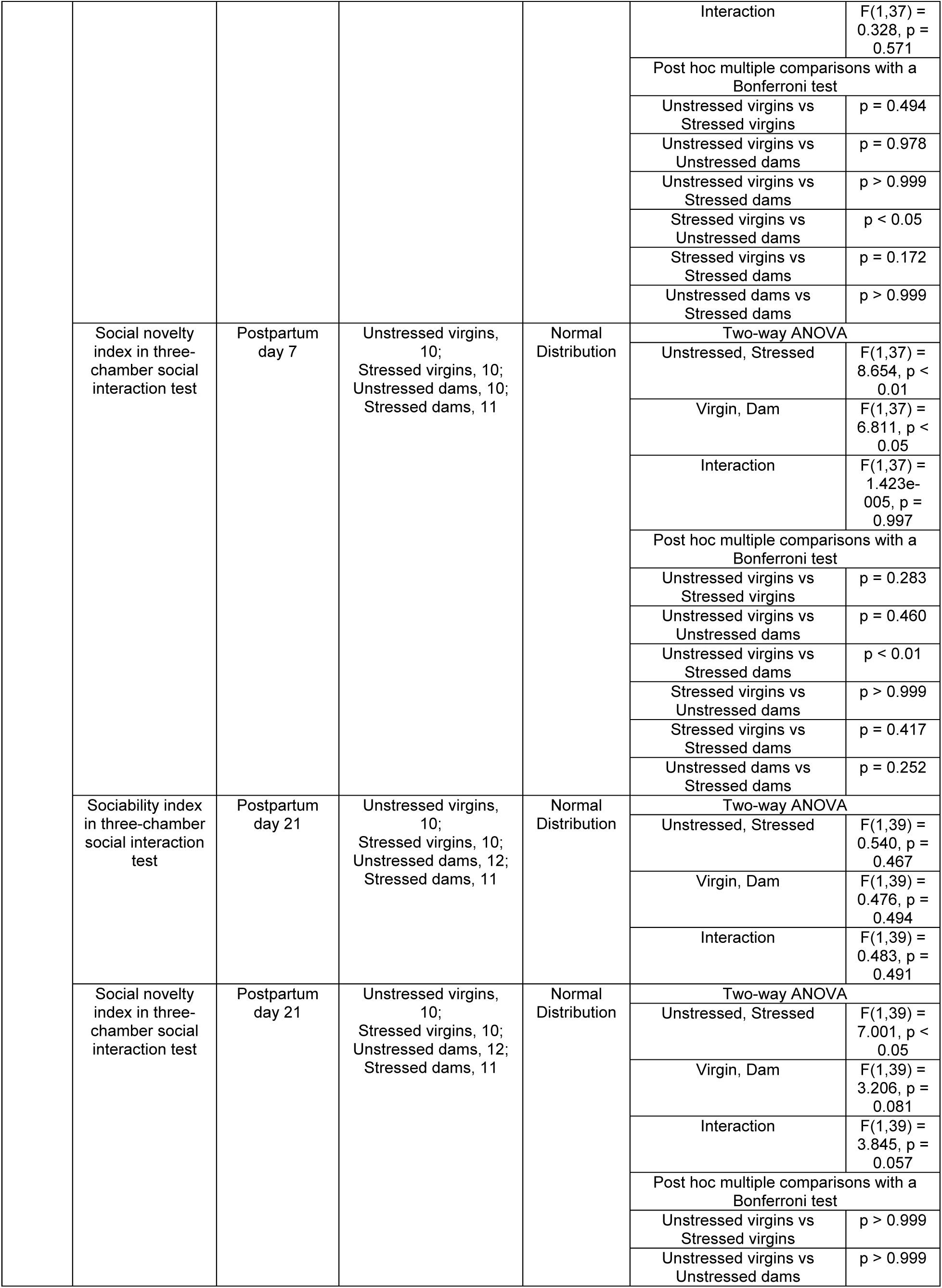
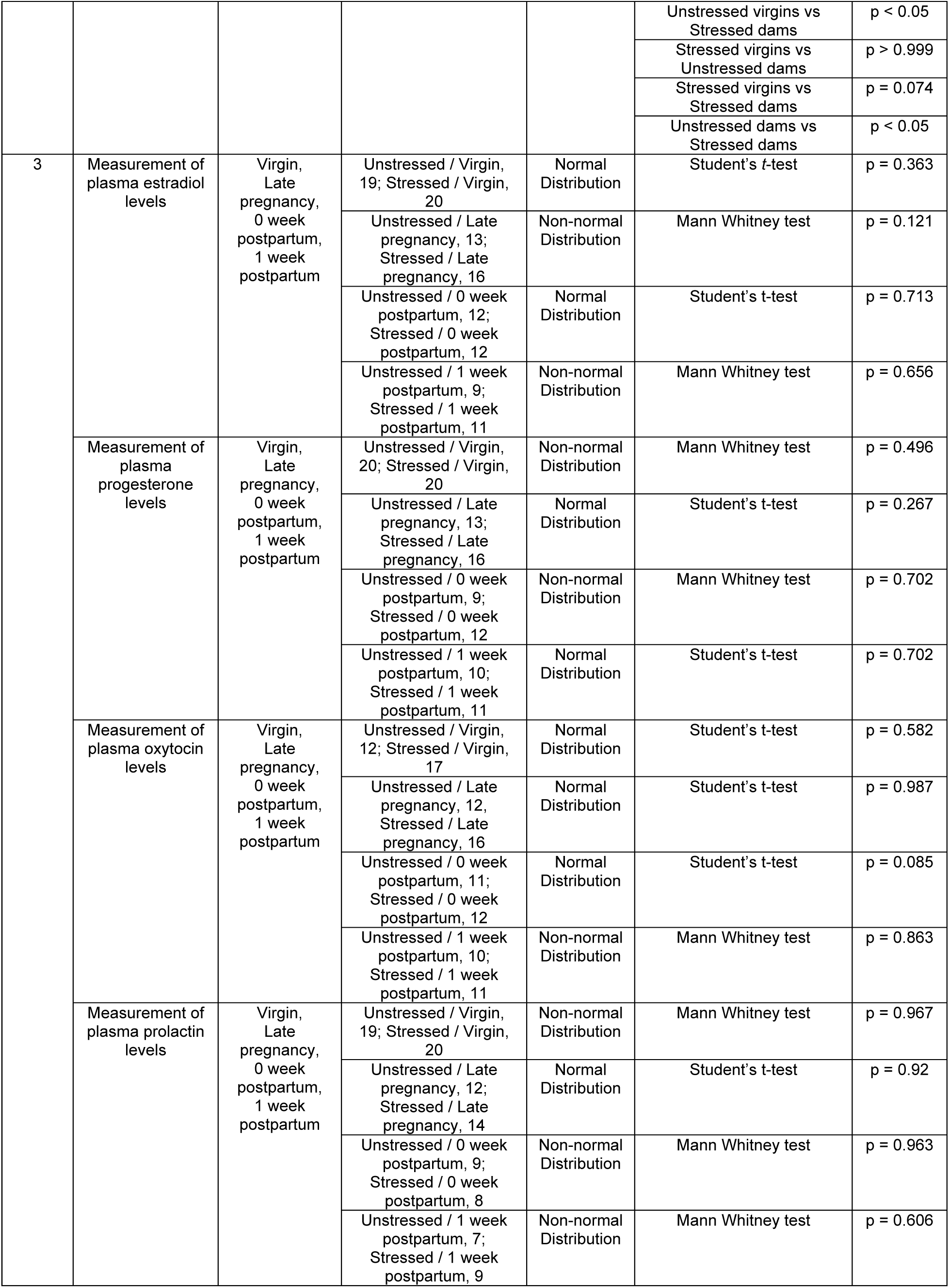
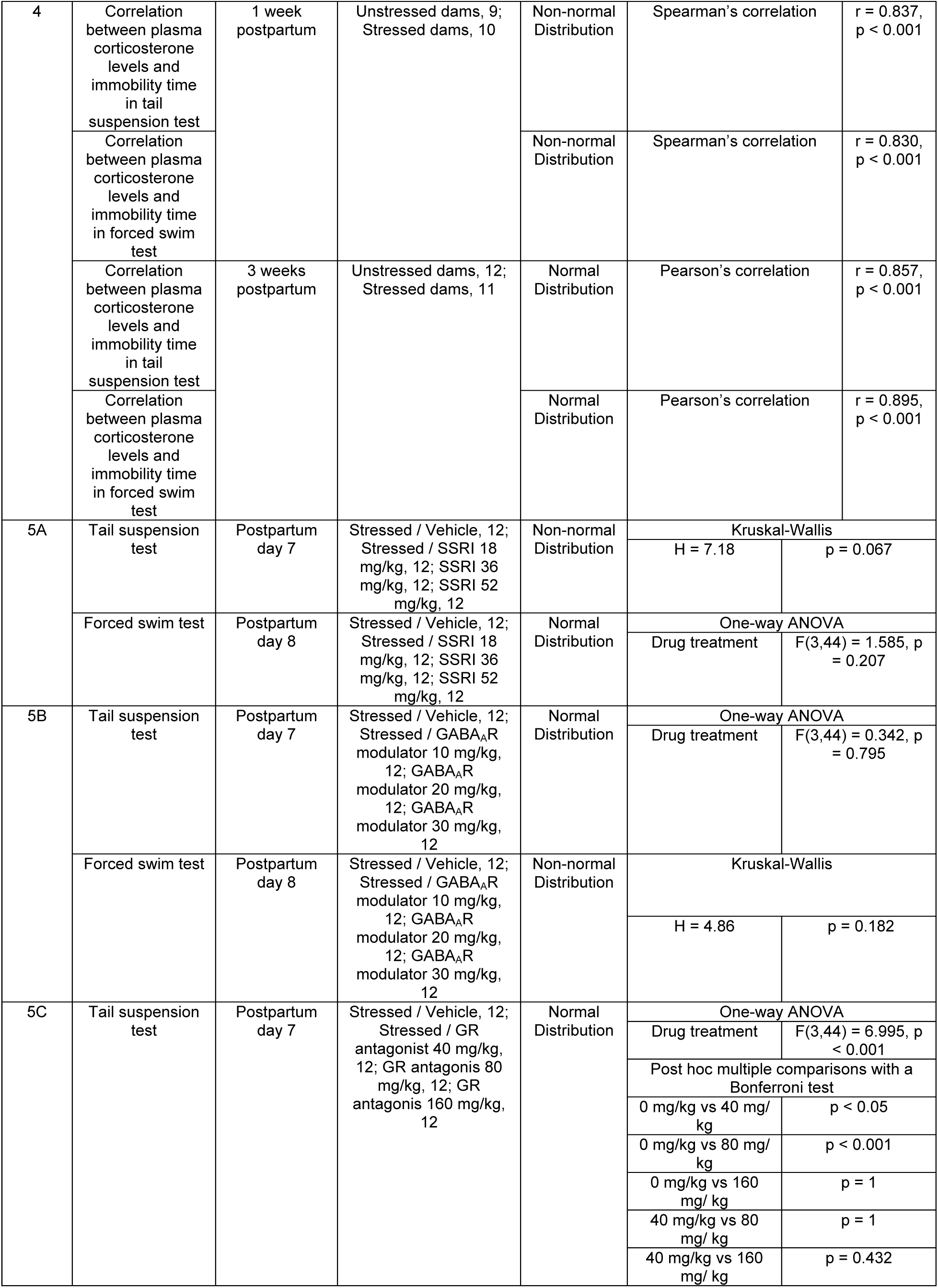
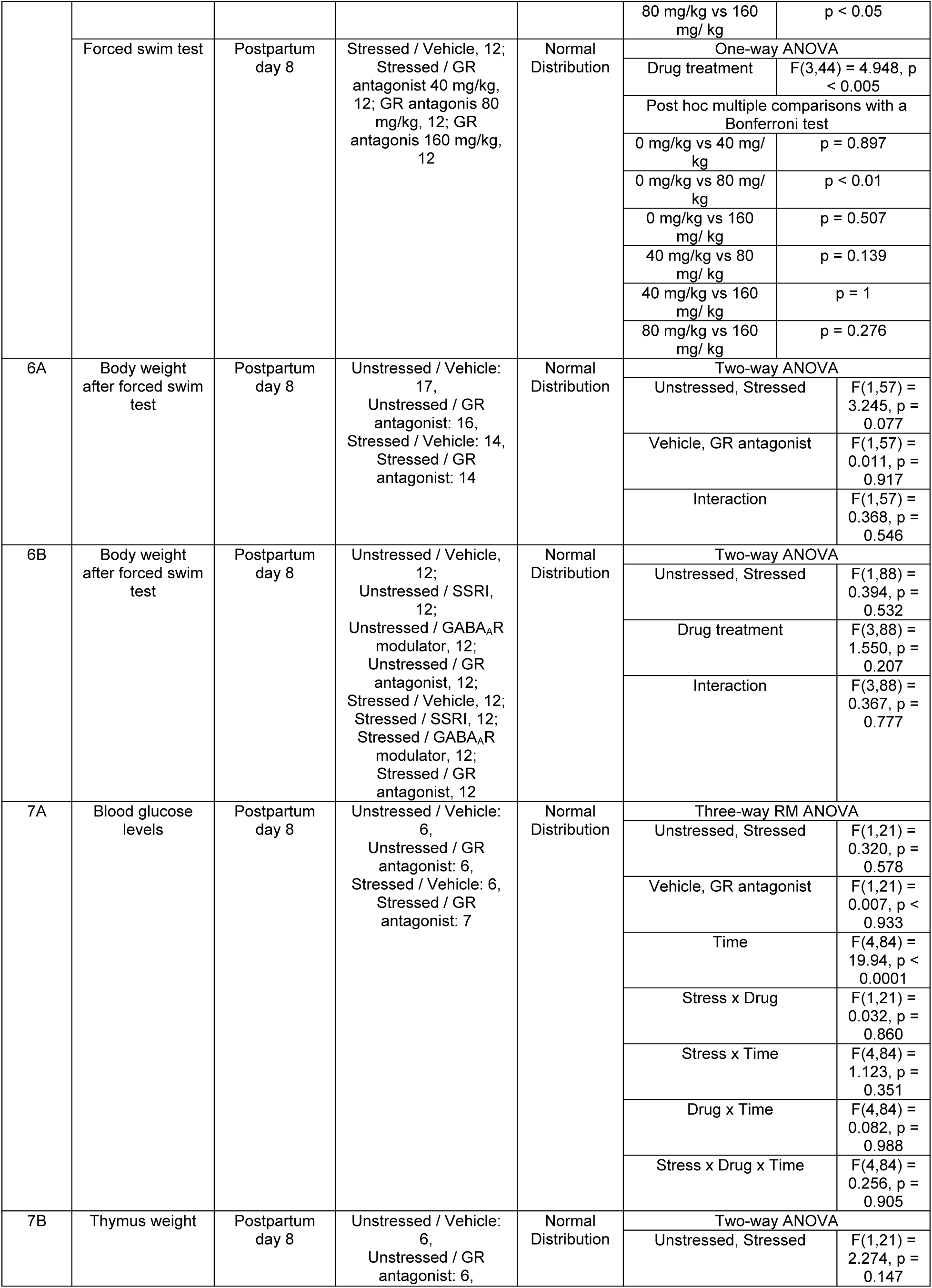
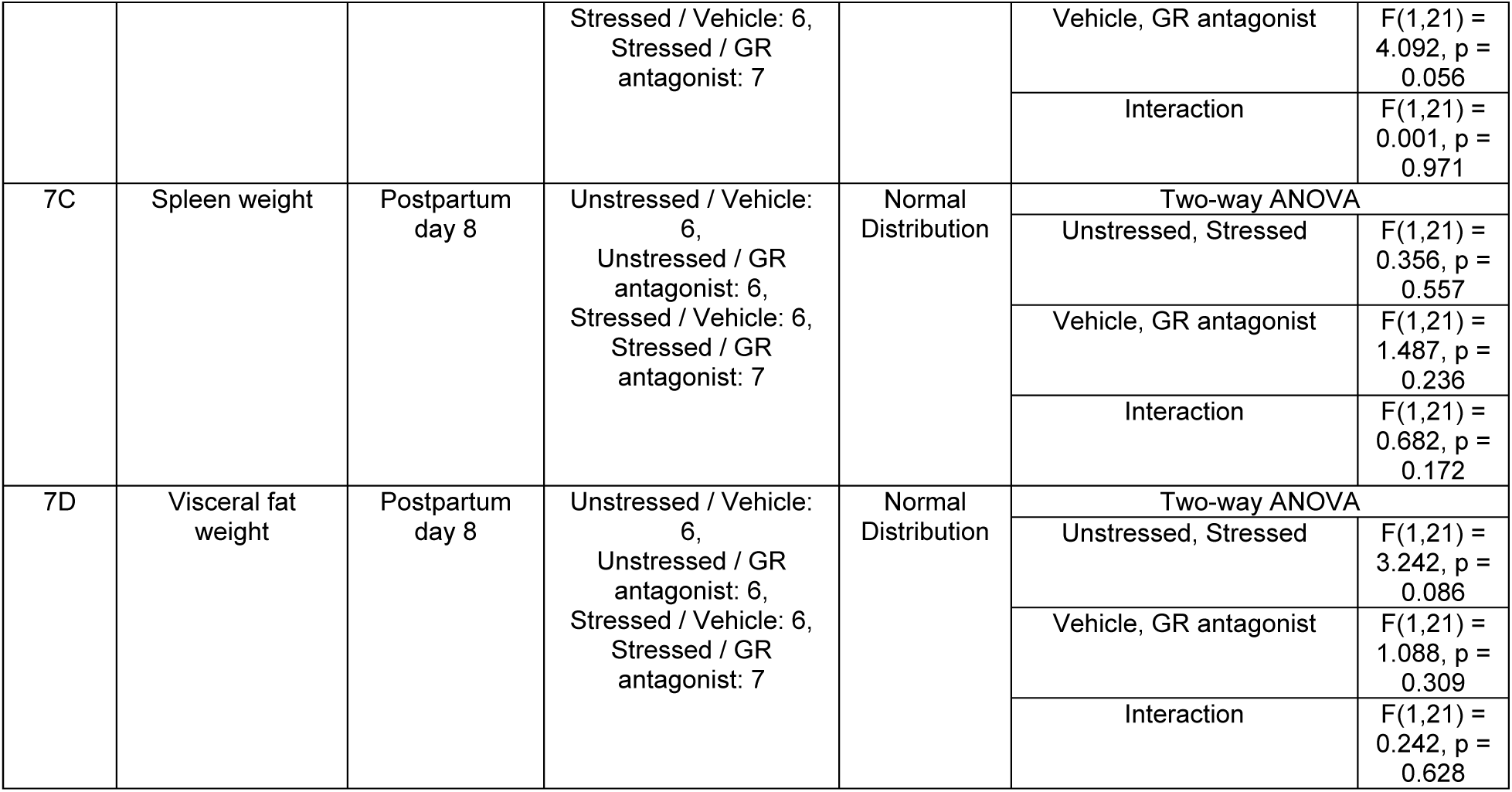
Statistical results.

## Notes

### Competing Interest Statement

The authors have declared no competing interest.

### Summary of Updates

Add two co-authors Add new data

## References

1 Brunton, P. J. & Russell, J. A. The expectant brain: adapting for motherhood. Nat Rev Neurosci 9, 11–25 (2008). 10.1038/nrn2280

2 O’Hara, M. W. & McCabe, J. E. Postpartum depression: current status and future directions. Annu Rev Clin Psychol 9, 379–407 (2013). 10.1146/annurev-clinpsy-050212-185612

3 Workman, J. L., Barha, C. K. & Galea, L. A. Endocrine substrates of cognitive and affective changes during pregnancy and postpartum. Behav Neurosci 126, 54–72 (2012). 10.1037/a0025538

4 Payne, J. L. & Maguire, J. Pathophysiological mechanisms implicated in postpartum depression. Front Neuroendocrinol 52, 165–180 (2019). 10.1016/j.yfrne.2018.12.001

5 Viguera, A. C. et al. Episodes of mood disorders in 2,252 pregnancies and postpartum periods. Am J Psychiatry 168, 1179–1185 (2011). 10.1176/appi.ajp.2011.11010148

6 Guintivano, J. et al. Adverse life events, psychiatric history, and biological predictors of postpartum depression in an ethnically diverse sample of postpartum women. Psychol Med 48, 1190–1200 (2018). 10.1017/S0033291717002641

7 Williams, L. M., Debattista, C., Duchemin, A. M., Schatzberg, A. F. & Nemeroff, C. B. Childhood trauma predicts antidepressant response in adults with major depression: data from the randomized international study to predict optimized treatment for depression. Transl Psychiatry 6, e799 (2016). 10.1038/tp.2016.61

8 Nemeroff, C. B. et al. Differential responses to psychotherapy versus pharmacotherapy in patients with chronic forms of major depression and childhood trauma. Proc Natl Acad Sci U S A 100, 14293–14296 (2003). 10.1073/pnas.2336126100

9 Schiller, C. E., Meltzer-Brody, S. & Rubinow, D. R. The role of reproductive hormones in postpartum depression. CNS Spectr 20, 48–59 (2015). 10.1017/S1092852914000480

10 Axelrod, J. & Reisine, T. D. Stress hormones: their interaction and regulation. Science 224, 452–459 (1984).

11 Joels, M. & Baram, T. Z. The neuro-symphony of stress. Nat Rev Neurosci 10, 459–466 (2009). 10.1038/nrn2632

12 Sorrells, S. F., Caso, J. R., Munhoz, C. D. & Sapolsky, R. M. The stressed CNS: when glucocorticoids aggravate inflammation. Neuron 64, 33–39 (2009). 10.1016/j.neuron.2009.09.032

13 Brummelte, S. & Galea, L. A. Postpartum depression: Etiology, treatment and consequences for maternal care. Horm Behav 77, 153–166 (2016). 10.1016/j.yhbeh.2015.08.008

14 Guintivano, J., Arad, M., Gould, T. D., Payne, J. L. & Kaminsky, Z. A. Antenatal prediction of postpartum depression with blood DNA methylation biomarkers. Mol Psychiatry 19, 560–567 (2014). 10.1038/mp.2013.62

15 Osborne, L. et al. Replication of Epigenetic Postpartum Depression Biomarkers and Variation with Hormone Levels. Neuropsychopharmacology 41, 1648–1658 (2016). 10.1038/npp.2015.333

16 Meltzer-Brody, S. et al. Brexanolone injection in post-partum depression: two multicentre, double-blind, randomised, placebo-controlled, phase 3 trials. Lancet 392, 1058–1070 (2018). 10.1016/S0140-6736(18)31551-4

17 Bloch, M., Daly, R. C. & Rubinow, D. R. Endocrine factors in the etiology of postpartum depression. Compr Psychiatry 44, 234–246 (2003). 10.1016/S0010-440X(03)00034-8

18 Ahokas, A., Kaukoranta, J., Wahlbeck, K. & Aito, M. Estrogen deficiency in severe postpartum depression: successful treatment with sublingual physiologic 17beta-estradiol: a preliminary study. J Clin Psychiatry 62, 332–336 (2001). 10.4088/jcp.v62n0504

19 Bloch, M. et al. Effects of gonadal steroids in women with a history of postpartum depression. Am J Psychiatry 157, 924–930 (2000). 10.1176/appi.ajp.157.6.924

20 Gregoire, A. J., Kumar, R., Everitt, B., Henderson, A. F. & Studd, J. W. Transdermal oestrogen for treatment of severe postnatal depression. Lancet 347, 930–933 (1996). 10.1016/s0140-6736(96)91414-2

21 Sichel, D. A., Cohen, L. S., Robertson, L. M., Ruttenberg, A. & Rosenbaum, J. F. Prophylactic estrogen in recurrent postpartum affective disorder. Biol Psychiatry 38, 814–818 (1995). 10.1016/0006-3223(95)00063-1

22 Galea, L. A., Wide, J. K. & Barr, A. M. Estradiol alleviates depressive-like symptoms in a novel animal model of post-partum depression. Behav Brain Res 122, 1–9 (2001). 10.1016/s0166-4328(01)00170-x

23 Green, A. D., Barr, A. M. & Galea, L. A. Role of estradiol withdrawal in ’anhedonic’ sucrose consumption: a model of postpartum depression. Physiol Behav 97, 259–265 (2009). 10.1016/j.physbeh.2009.02.020

24 Zhang, Z. et al. Postpartum estrogen withdrawal impairs hippocampal neurogenesis and causes depression- and anxiety-like behaviors in mice. Psychoneuroendocrinology 66, 138–149 (2016). 10.1016/j.psyneuen.2016.01.013

25 Stewart, D. E. & Vigod, S. N. Postpartum Depression: Pathophysiology, Treatment, and Emerging Therapeutics. Annu Rev Med 70, 183–196 (2019). 10.1146/annurev-med-041217-011106

26 Molyneaux, E., Trevillion, K. & Howard, L. M. Antidepressant treatment for postnatal depression. JAMA 313, 1965–1966 (2015). 10.1001/jama.2015.2276

27 Langan, R. & Goodbred, A. J. Identification and Management of Peripartum Depression. Am Fam Physician 93, 852–858 (2016).

28 Meltzer-Brody, S. & Kanes, S. J. Allopregnanolone in postpartum depression: Role in pathophysiology and treatment. Neurobiol Stress 12, 100212 (2020). 10.1016/j.ynstr.2020.100212

29 Kanes, S. et al. Brexanolone (SAGE-547 injection) in post-partum depression: a randomised controlled trial. Lancet 390, 480–489 (2017). 10.1016/S0140-6736(17)31264-3

30 Leader, L. D., O’Connell, M. & VandenBerg, A. Brexanolone for Postpartum Depression: Clinical Evidence and Practical Considerations. Pharmacotherapy 39, 1105–1112 (2019). 10.1002/phar.2331

31 Blakemore, S. J. The social brain in adolescence. Nat Rev Neurosci 9, 267–277 (2008). 10.1038/nrn2353

32 Ibi, D. et al. Social isolation rearing-induced impairment of the hippocampal neurogenesis is associated with deficits in spatial memory and emotion-related behaviors in juvenile mice. J Neurochem 105, 921–932 (2008). 10.1111/j.1471-4159.2007.05207.x

33 Kin, K., Gaini, R. & Niwa, M. in Encyclopedia of Behavioral Neuroscience*, Second Edition* Vol. 1 (ed Sergio Della Sala) 360–371 (Elsevier, 2021).

34 Peters, Y. M. & O’Donnell, P. Social isolation rearing affects prefrontal cortical response to ventral tegmental area stimulation. Biol Psychiatry 57, 1205–1208 (2005). 10.1016/j.biopsych.2005.02.011

35 Lukkes, J. L., Watt, M. J., Lowry, C. A. & Forster, G. L. Consequences of post-weaning social isolation on anxiety behavior and related neural circuits in rodents. Front Behav Neurosci 3, 18 (2009). 10.3389/neuro.08.018.2009

36 Walker, D. M., Cunningham, A. M., Gregory, J. K. & Nestler, E. J. Long-Term Behavioral Effects of Post-weaning Social Isolation in Males and Females. Front Behav Neurosci 13, 66 (2019). 10.3389/fnbeh.2019.00066

37 Niwa, M. et al. Adolescent stress-induced epigenetic control of dopaminergic neurons via glucocorticoids. Science 339, 335–339 (2013). 10.1126/science.1226931

38 Niwa, M. et al. A critical period of vulnerability to adolescent stress: epigenetic mediators in mesocortical dopaminergic neurons. Hum Mol Genet 25, 1370–1381 (2016). 10.1093/hmg/ddw019

39 Hikida, T. et al. Adolescent psychosocial stress enhances sensitization to cocaine exposure in genetically vulnerable mice. Neurosci Res 151, 38–45 (2020). 10.1016/j.neures.2019.02.007

40 Matsumoto, Y. et al. Adolescent stress leads to glutamatergic disturbance through dopaminergic abnormalities in the prefrontal cortex of genetically vulnerable mice. Psychopharmacology (Berl) 234, 3055–3074 (2017). 10.1007/s00213-017-4704-8

41 Kin, K., Francis-Oliveira, J., Kano, S. I. & Niwa, M. Adolescent stress impairs postpartum social behavior via anterior insula-prelimbic pathway in mice. Nat Commun 14, 2975 (2023). 10.1038/s41467-023-38799-6

42 Molendijk, M. L. & de Kloet, E. R. Immobility in the forced swim test is adaptive and does not reflect depression. Psychoneuroendocrinology 62, 389–391 (2015). 10.1016/j.psyneuen.2015.08.028

43 Melon, L., Hammond, R., Lewis, M. & Maguire, J. A Novel, Synthetic, Neuroactive Steroid Is Effective at Decreasing Depression-Like Behaviors and Improving Maternal Care in Preclinical Models of Postpartum Depression. Front Endocrinol (Lausanne) 9, 703 (2018). 10.3389/fendo.2018.00703

44 Liu, M. Y. et al. Sucrose preference test for measurement of stress-induced anhedonia in mice. Nat Protoc 13, 1686–1698 (2018). 10.1038/s41596-018-0011-z

45 Millan, M. J. & Bales, K. L. Towards improved animal models for evaluating social cognition and its disruption in schizophrenia: the CNTRICS initiative. Neurosci Biobehav Rev 37, 2166–2180 (2013). 10.1016/j.neubiorev.2013.09.012

46 Yang, M., Silverman, J. L. & Crawley, J. N. Automated three-chambered social approach task for mice. Curr Protoc Neurosci **Chapter** 8, Unit 8 26 (2011). 10.1002/0471142301.ns0826s56

47 Hendrick, V., Altshuler, L. L. & Suri, R. Hormonal changes in the postpartum and implications for postpartum depression. Psychosomatics 39, 93–101 (1998). 10.1016/S0033-3182(98)71355-6

48 Stoffel, E. C. & Craft, R. M. Ovarian hormone withdrawal-induced "depression" in female rats. Physiol Behav 83, 505–513 (2004). 10.1016/j.physbeh.2004.08.033

49 Suda, S., Segi-Nishida, E., Newton, S. S. & Duman, R. S. A postpartum model in rat: behavioral and gene expression changes induced by ovarian steroid deprivation. Biol Psychiatry 64, 311–319 (2008). 10.1016/j.biopsych.2008.03.029

50 Lupien, S. J., McEwen, B. S., Gunnar, M. R. & Heim, C. Effects of stress throughout the lifespan on the brain, behaviour and cognition. Nat Rev Neurosci 10, 434–445 (2009). 10.1038/nrn2639

51 Herman, J. P., Nawreen, N., Smail, M. A. & Cotella, E. M. Brain mechanisms of HPA axis regulation: neurocircuitry and feedback in context Richard Kvetnansky lecture. Stress 23, 617–632 (2020). 10.1080/10253890.2020.1859475

52 De Kloet, E. R. Why Dexamethasone Poorly Penetrates in Brain. Stress 2, 13–20 (1997).

53 Meijer, O. C. et al. Penetration of dexamethasone into brain glucocorticoid targets is enhanced in mdr1A P-glycoprotein knockout mice. Endocrinology 139, 1789–1793 (1998). 10.1210/endo.139.4.5917

54 Liston, C. & Gan, W. B. Glucocorticoids are critical regulators of dendritic spine development and plasticity in vivo. Proc Natl Acad Sci U S A 108, 16074–16079 (2011). 10.1073/pnas.1110444108

55 De Kloet, E. R., Vreugdenhil, E., Oitzl, M. S. & Joels, M. Brain corticosteroid receptor balance in health and disease. Endocr Rev 19, 269–301 (1998). 10.1210/edrv.19.3.0331

56 McEwen, B. S. Physiology and neurobiology of stress and adaptation: central role of the brain. Physiol Rev 87, 873–904 (2007). 10.1152/physrev.00041.2006

57 Ogino, S., Fuchs, C. S. & Giovannucci, E. How many molecular subtypes? Implications of the unique tumor principle in personalized medicine. Expert Rev Mol Diagn 12, 621–628 (2012). 10.1586/erm.12.46

58 Waks, A. G. & Winer, E. P. Breast Cancer Treatment: A Review. JAMA 321, 288–300 (2019). 10.1001/jama.2018.19323

59 Collisson, E. A., Bailey, P., Chang, D. K. & Biankin, A. V. Molecular subtypes of pancreatic cancer. Nat Rev Gastroenterol Hepatol 16, 207–220 (2019). 10.1038/s41575-019-0109-y

60 Walton, N. & Maguire, J. Allopregnanolone-based treatments for postpartum depression: Why/how do they work? Neurobiol Stress 11, 100198 (2019). 10.1016/j.ynstr.2019.100198

61 Mody, I. GABAAR Modulator for Postpartum Depression. Cell 176, 1 (2019). 10.1016/j.cell.2018.12.016

62 Belelli, D., Hogenkamp, D., Gee, K. W. & Lambert, J. J. Realising the therapeutic potential of neuroactive steroid modulators of the GABAA receptor. Neurobiol Stress 12, 100207 (2020). 10.1016/j.ynstr.2019.100207

63 Althaus, A. L. et al. Preclinical characterization of zuranolone (SAGE-217), a selective neuroactive steroid GABAA receptor positive allosteric modulator. Neuropharmacology 181, 108333 (2020). 10.1016/j.neuropharm.2020.108333

64 Beaudry, J. L. et al. Effects of selective and non-selective glucocorticoid receptor II antagonists on rapid-onset diabetes in young rats. PLoS One 9, e91248 (2014). 10.1371/journal.pone.0091248

65 Asagami, T. et al. Selective Glucocorticoid Receptor (GR-II) Antagonist Reduces Body Weight Gain in Mice. J Nutr Metab 2011, 235389 (2011). 10.1155/2011/235389

66 Samuels, B. A. et al. 5-HT1A receptors on mature dentate gyrus granule cells are critical for the antidepressant response. Nat Neurosci 18, 1606–1616 (2015). 10.1038/nn.4116

67 Locci, A., Geoffroy, P., Miesch, M., Mensah-Nyagan, A. G. & Pinna, G. Social Isolation in Early versus Late Adolescent Mice Is Associated with Persistent Behavioral Deficits That Can Be Improved by Neurosteroid-Based Treatment. Front Cell Neurosci 11, 208 (2017). 10.3389/fncel.2017.00208

68 Gehrand, A. L. et al. Glucocorticoid Receptor Antagonist Alters Corticosterone and Receptor-sensitive mRNAs in the Hypoxic Neonatal Rat. Endocrinology 163 (2022). 10.1210/endocr/bqab232

69 Kroon, J. et al. Selective Glucocorticoid Receptor Antagonist CORT125281 Activates Brown Adipose Tissue and Alters Lipid Distribution in Male Mice. Endocrinology 159, 535–546 (2018). 10.1210/en.2017-00512

70 McEwen, B. S. & Morrison, J. H. The brain on stress: vulnerability and plasticity of the prefrontal cortex over the life course. Neuron 79, 16–29 (2013). 10.1016/j.neuron.2013.06.028

71 McEwen, B. S. & Gianaros, P. J. Central role of the brain in stress and adaptation: links to socioeconomic status, health, and disease. Ann N Y Acad Sci 1186, 190–222 (2010). 10.1111/j.1749-6632.2009.05331.x

72 Arnsten, A. F. Stress signalling pathways that impair prefrontal cortex structure and function. Nat Rev Neurosci 10, 410–422 (2009). 10.1038/nrn2648

73 Craig, A. D. How do you feel--now? The anterior insula and human awareness. Nat Rev Neurosci 10, 59–70 (2009). 10.1038/nrn2555

74 Critchley, H. D. & Harrison, N. A. Visceral influences on brain and behavior. Neuron 77, 624–638 (2013). 10.1016/j.neuron.2013.02.008

75 Nestler, E. J. & Hyman, S. E. Animal models of neuropsychiatric disorders. Nat Neurosci 13, 1161–1169 (2010). 10.1038/nn.2647

76 Perlman, R. L. Mouse models of human disease: An evolutionary perspective. Evol Med Public Health 2016, 170–176 (2016). 10.1093/emph/eow014

77 Castinetti, F., Brue, T. & Conte-Devolx, B. The use of the glucocorticoid receptor antagonist mifepristone in Cushing’s syndrome. Curr Opin Endocrinol Diabetes Obes 19, 295–299 (2012). 10.1097/MED.0b013e32835430bf

78 Hunt, H. et al. Assessment of Safety, Tolerability, Pharmacokinetics, and Pharmacological Effect of Orally Administered CORT125134: An Adaptive, Double-Blind, Randomized, Placebo-Controlled Phase 1 Clinical Study. Clin Pharmacol Drug Dev 7, 408–421 (2018). 10.1002/cpdd.389

79 Hunt, H. J. et al. Identification of the Clinical Candidate (R)-(1-(4-Fluorophenyl)-6-((1-methyl-1H-pyrazol-4-yl)sulfonyl)-4,4a,5,6,7,8-hexah ydro-1H-pyrazolo[3,4-g]isoquinolin-4a-yl)(4-(trifluoromethyl)pyridin-2-yl)methano ne (CORT125134): A Selective Glucocorticoid Receptor (GR) Antagonist. J Med Chem 60, 3405–3421 (2017). 10.1021/acs.jmedchem.7b00162

80 Vendruscolo, L. F. et al. Glucocorticoid receptor antagonism decreases alcohol seeking in alcohol-dependent individuals. J Clin Invest 125, 3193–3197 (2015). 10.1172/Jci79828

81 Pineau, F. et al. New selective glucocorticoid receptor modulators reverse amyloid-beta peptide-induced hippocampus toxicity. Neurobiol Aging 45, 109–122 (2016). 10.1016/j.neurobiolaging.2016.05.018

82 Hunt, H. J. et al. 1H-Pyrazolo[3,4-g]hexahydro-isoquinolines as potent GR antagonists with reduced hERG inhibition and an improved pharmacokinetic profile. Bioorg Med Chem Lett 25, 5720–5725 (2015). 10.1016/j.bmcl.2015.10.097

83 Dulawa, S. C., Holick, K. A., Gundersen, B. & Hen, R. Effects of chronic fluoxetine in animal models of anxiety and depression. Neuropsychopharmacology 29, 1321–1330 (2004). 10.1038/sj.npp.1300433

84 McMurray, K. M. J. et al. Identification of a novel, fast-acting GABAergic antidepressant. Mol Psychiatry 23, 384–391 (2018). 10.1038/mp.2017.14

85 Siopi, E. et al. Anxiety- and Depression-Like States Lead to Pronounced Olfactory Deficits and Impaired Adult Neurogenesis in Mice. J Neurosci 36, 518–531 (2016). 10.1523/JNEUROSCI.2817-15.2016

86 Amellem, I., Suresh, S., Chang, C. C., Tok, S. S. L. & Tashiro, A. A critical period for antidepressant-induced acceleration of neuronal maturation in adult dentate gyrus. Transl Psychiatry 7, e1235 (2017). 10.1038/tp.2017.208

87 Turcotte-Cardin, V. et al. Loss of Adult 5-HT1A Autoreceptors Results in a Paradoxical Anxiogenic Response to Antidepressant Treatment. J Neurosci 39, 1334–1346 (2019). 10.1523/JNEUROSCI.0352-18.2018

88 Vahid-Ansari, F. et al. Abrogated Freud-1/Cc2d1a Repression of 5-HT1A Autoreceptors Induces Fluoxetine-Resistant Anxiety/Depression-Like Behavior. J Neurosci 37, 11967–11978 (2017). 10.1523/JNEUROSCI.1668-17.2017

89 Kazdoba, T. M., Hagerman, R. J., Zolkowska, D., Rogawski, M. A. & Crawley, J. N. Evaluation of the neuroactive steroid ganaxolone on social and repetitive behaviors in the BTBR mouse model of autism. Psychopharmacology (Berl) 233, 309–323 (2016). 10.1007/s00213-015-4115-7

90 Savarese, A. M. et al. Targeting the Glucocorticoid Receptor Reduces Binge-Like Drinking in High Drinking in the Dark (HDID-1) Mice. Alcohol Clin Exp Res 44, 1025–1036 (2020). 10.1111/acer.14318

91 Pinna, G. & Rasmusson, A. M. Ganaxolone improves behavioral deficits in a mouse model of post-traumatic stress disorder. Front Cell Neurosci 8, 256 (2014). 10.3389/fncel.2014.00256

92 Cao, X. et al. Astrocyte-derived ATP modulates depressive-like behaviors. Nat Med 19, 773–777 (2013). 10.1038/nm.3162

93 Rein, B., Ma, K. & Yan, Z. A standardized social preference protocol for measuring social deficits in mouse models of autism. Nat Protoc 15, 3464–3477 (2020). 10.1038/s41596-020-0382-9

94 Niwa, M. et al. Knockdown of DISC1 by in utero gene transfer disturbs postnatal dopaminergic maturation in the frontal cortex and leads to adult behavioral deficits. Neuron 65, 480–489 (2010). 10.1016/j.neuron.2010.01.019

95 Boyle, M. P. et al. Acquired deficit of forebrain glucocorticoid receptor produces depression-like changes in adrenal axis regulation and behavior. Proc Natl Acad Sci U S A 102, 473–478 (2005). 10.1073/pnas.0406458102

96 Pang, T. Y. et al. Positive environmental modification of depressive phenotype and abnormal hypothalamic-pituitary-adrenal axis activity in female C57BL/6J mice during abstinence from chronic ethanol consumption. Front Pharmacol 4, 93 (2013). 10.3389/fphar.2013.00093

97 Andrikopoulos, S., Blair, A. R., Deluca, N., Fam, B. C. & Proietto, J. Evaluating the glucose tolerance test in mice. Am J Physiol Endocrinol Metab 295, E1323–1332 (2008). 10.1152/ajpendo.90617.2008

98 Bowe, J. E. et al. Metabolic phenotyping guidelines: assessing glucose homeostasis in rodent models. J Endocrinol 222, G13–25 (2014). 10.1530/JOE-14-0182

99 Kimmel, M. et al. Family history, not lack of medication use, is associated with the development of postpartum depression in a high-risk sample. Arch Womens Ment Health 18, 113–121 (2015). 10.1007/s00737-014-0432-9

